# Benchmarking of bulk transcriptomic harmonization tools in a multi-platform B-cell lymphoma cohort identifies feature-specific quantile normalization and surrogate variable analysis as top-performing methods

**DOI:** 10.64898/2026.09.15.751825

**Authors:** Daniil Nikitin, Nikolay Borisov, Maria Savchenko, Anatoly Bobe, Mark Meerson, Alexander Nesmelov, Nazar Harutyunyan, Svetlana Paponova, Andrey Kravets, Alexandr Zaitsev, Alexandr Bagaev, Arsen Arakelyan

## Abstract

Cross-platform harmonization of bulk transcriptomic datasets remains a fundamental challenge for developing cancer biomarkers because of persistent unresolved batch effects. Most harmonization tools are benchmarked on datasets with large inter-group biological differences (for example TCGA tumor types), whereas actionable biomarker mining requires preserving subtle transcriptional distinctions between closely related diagnoses. Here we present ComboBatch, a benchmarking pipeline that evaluates the full cross-product of 14 batch-removal strategies, 3 imputation methods, 33 harmonization algorithms and 2 post-removal conditions across 7,174 samples from 88 germinal-center B-cell lymphoma cohorts spanning four transcriptomic platforms. Scoring 87 quality metrics across 2,234 harmonization approaches, we show that method choice (R^2^ 0.36) and batch-removal strategy (0.26) are the principal determinants of harmonization quality, whereas imputation (0.016) and post-removal (<0.01) are secondary. Feature Specific Quantile Normalization and Surrogate Variable Analysis were the top methods, jointly resolving follicular lymphoma, diffuse large B-cell lymphoma and normal germinal-center B-cell differences in multi-platform and RNA-seq-only compositions, respectively. We provide a data-driven five-scenario decision tree for harmonization method selection, applicable to any retrospective multi-platform transcriptomic study. The ComboBatch pipeline is available on GitHub and can be used for harmonization, allowing bioinformaticians to utilize 33 harmonization and 3 imputation methods according to their needs.

**GRAPHICAL ABSTRACT:** 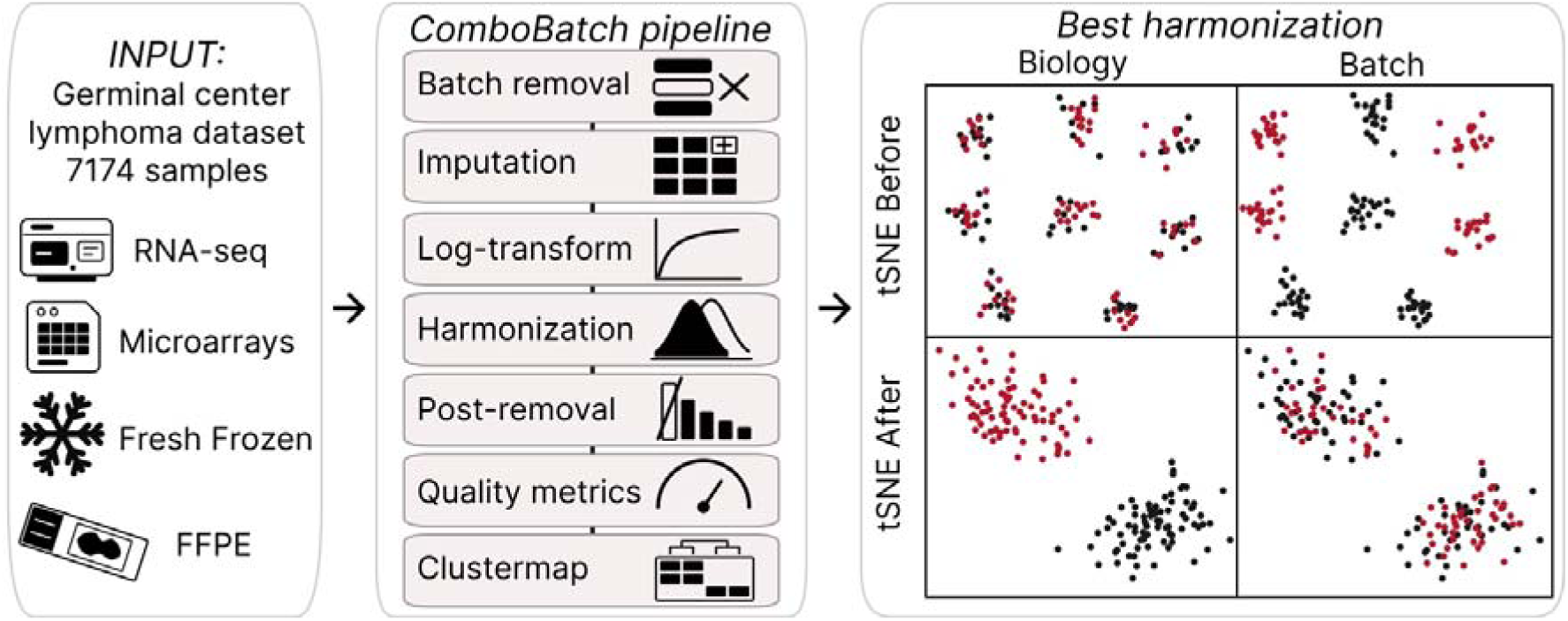

## 1. INTRODUCTION

Despite considerable progress in transcriptomic batch correction, technical variation introduced by differences in sequencing platform, RNA extraction protocol, and sample preservation method remains a principal obstacle to the development of reproducible cancer biomarkers (Vladimirova et al. 2021a; Gudkov et al. 2022a) — particularly for AI-driven classifiers trained on heterogeneous multi-cohort datasets (Yu et al. 2024). Batch-confounded AI models can undergo performance collapse upon clinical deployment, as exemplified by an area under the curve (AUC) decrease from 0.79 to 0.50 reported for a digital pathology prognostic classifier (Dawood et al. 2026 Mar 2).

Despite the rise of single-cell RNA sequencing, bulk transcriptomic profiling remains the standard approach for biomarker detection and therapy response prediction, justified by the fact that there are approximately 65,000 bulk-profiled patients (Zhang et al. 2021; Cho et al. 2026) compared with approximately 4,000 single-cell-profiled patients (∼ 10 million cells) (Liu et al. 2025; Zeng et al. 2025).

The two principal sources of batch variation in multi-platform bulk transcriptomic studies are the technology axis — microarray versus RNA-seq (Raplee et al. 2025) — and the biomaterial axis — fresh-frozen (FF) versus formalin-fixed paraffin-embedded (FFPE) tissue (Newton et al. 2020) — which introduce systematic differences in gene coverage, dynamic range, and signal-to-noise ratio (Yu et al. 2024).

Most harmonization tools have been developed and benchmarked on datasets characterized by large inter-group biological differences, such as TCGA pan-cancer collections (Yu et al. 2024), in which the biological signal substantially exceeds technical noise. However, the majority of cancer studies analyze related subtypes (Shtam et al. 2018; Jovčevska et al. 2019; Ma et al. 2024) — such as follicular lymphoma (FL) and diffuse large B-cell lymphoma (DLBCL), which share a common germinal center (GC) B-cell origin and exhibit partially overlapping transcriptional programs (Laurent et al. 2024). These studies operate in a fundamentally different signal-to-noise regime, in which technical batch effects can substantially exceed the biology of interest (Dai et al. 2025).

To benchmark the available harmonization tools under conditions of subtle biological and large-scale technical variation, we assembled a dataset of 7,174 DLBCL, FL and normal B-cell samples. The dataset consisted of 88 individual cohorts, profiled under RNA-seq as well as Affymetrix, Illumina and Agilent microarrays and encompassing FFPE, FF and other biomaterial types (e.g. sorted cells). Batch-explained variance was ten times higher than the biology-explained variance according to Principal Component Regression estimate (PCReg).

We built a computational pipeline, ComboBatch, that comprised 14 strategies of prior batch removal, 3 imputation methods, 31 harmonization methods and an optional post harmonization removal step – encompassing 2,234 successfully completed harmonization approaches. We then utilized 87 quality metrics and showed that method choice (R^2^ 0.36) and batch-removal strategy (0.26) are the principal determinants of harmonization quality, whereas imputation (0.016) and post-removal (<0.01) are secondary.

We demonstrated that Feature Specific Quantile Normalization (FSQN) and Surrogate Variable Analysis (SVA) were the top methods, jointly resolving FL, DLBCL and normal B-cell differences in multi-platform and RNA-seq-only compositions, respectively. These methods were both the best performing ones by metrics and mixed batch classes together in t-distributed stochastic neighbor embedding (tSNE) and uniform manifold approximation and projection (UMAP) space, while preserving biology groups as distinct clouds and biomarker genes correlation before and after harmonization. We propose a data-driven five-scenario decision tree for harmonization method selection, applicable to any retrospective multi-platform transcriptomic study.

Finally, the ComboBatch tool is available in a public GitHub repository via the link https://github.com/Nikit357/ComboBatch. The tool includes code for evaluating 33 harmonization and 3 imputation methods with reusable Python environment for any bulk transcriptomic dataset. A Docker image can be deployed and integrated in any bioinformatic pipeline for harmonization if necessary.

Together, the benchmark and the ComboBatch tool provide a basis for harmonizing retrospective multi-platform transcriptomic datasets for biomarker discovery.

## 2. MATERIALS AND METHODS

### 2.1. Dataset assembly

We assembled a comprehensive set of publicly available cohorts containing FL samples, supplemented with DLBCL cohorts as the most closely related GC B-cell malignancy and normal B-cell cohorts as a biological reference for future FL transcriptomic signature studies. The final dataset comprised 88 cohorts (14 of them proprietary, the rest cohorts are open source), 7,174 samples, and four platform types: RNA-seq (NGS), Affymetrix, Illumina, and Agilent microarrays (Supplementary File 1). This study analyzed only publicly available, de-identified transcriptomic data and generated no new human data, so no additional ethics approval was required.

### 2.2. Batch annotation and classification

We devised a systematic batch naming convention in which each batch is labeled <PLATFORM_NAME>_<BIOMATERIAL_NAME>_<TARGET_ENRICHMENT_NAME>. Platform_name is the Gene Expression Omnibus (GEO) platform identifier for microarray batches, or ’RNASeq’ for NGS batches. Biomaterial_name is coded as ’FFPE’ for cohorts with unambiguous FFPE annotation, ’FF’ for all fresh biomaterials (fresh-frozen tissue, sorted cells, peripheral blood mononuclear cells (PBMCs), whole blood, etc.), and ’Unknown’ for samples lacking unambiguous biomaterial classification in the source article or GEO record. For NGS batches, Target_enrichment_name records the enrichment strategy (poly-A selection, exome capture, rRNA depletion, total RNA, or unknown); for microarray batches this field is set to ’unknown’, as target specificity is inherent to the probe design of the array platform. Additional granular batch attributes — RNA extraction kit, library preparation protocol, NGS platform model, and detailed biomaterial subtype — are available in Supplementary File 1 and in the GEO records. We excluded these fields from the batch classification scheme to avoid over-parameterizing the batch correction logic.

### 2.3. Gene expression profiling

For RNA-seq datasets gene expression profiling was done starting from the raw FASTQ files using the GRCh38.d1.vd1 reference, kallisto version 0.43.0 and GENCODE release 23 as described in (Kotlov et al. 2024). For microarray datasets, raw probe-level gene expression values were obtained from the GEO repository for each dataset’s corresponding platform. Probe identifiers were mapped to standard HGNC gene symbols using the Python biomart package (version 0.9.2) (GitHub - sebriois/biomart: Python biomart API · GitHub) querying the Ensembl BioMart database (release 116). For probes with no corresponding HGNC symbol, or corresponding to withdrawn/deprecated identifiers, entries were excluded from downstream analysis. For genes targeted by multiple probes, expression values were log-transformed using the formula log_2_(expression + 1) and the log-transformed values were averaged across probes to obtain a single gene-level expression value per sample. After the averaging, the expression values were back-transformed using the formula 2^expression^ – 1.

### 2.4. Computational hardware

The full cross-product of prior batch removal, imputation, log2-transformation, harmonization, and post-removal was executed on Kubernetes pods with 32–48 CPU cores and 240 GiB RAM, provisioned by the Karpenter node autoscaler (GitHub - kubernetes-sigs/karpenter: Karpenter is a Kubernetes Node Autoscaler built for flexibility, performance, and simplicity. · GitHub). Downstream data analysis and visualization were performed on a JupyterHub server with 16 CPU cores and 64 GiB RAM.

### 2.5. Batch removal strategies

Because most harmonization algorithms process samples jointly and their performance depends critically on the initial batch composition of the input dataset (Tzec-Interián et al. 2025), we designed 14 prior batch removal strategies to be applied before imputation and log2-transformation. These strategies were developed iteratively across three rounds of pipeline refinement (Table 1).

**Table 1.**
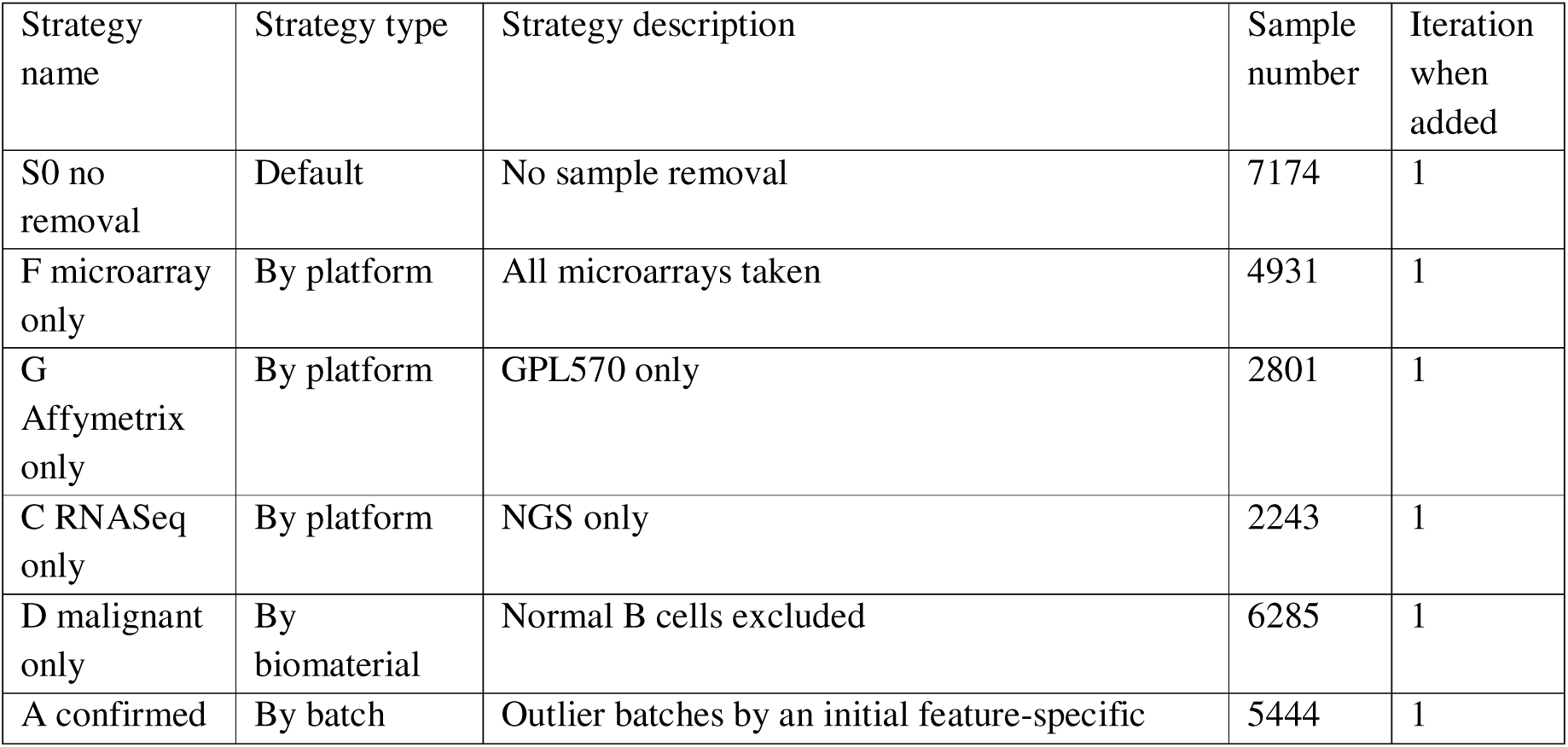
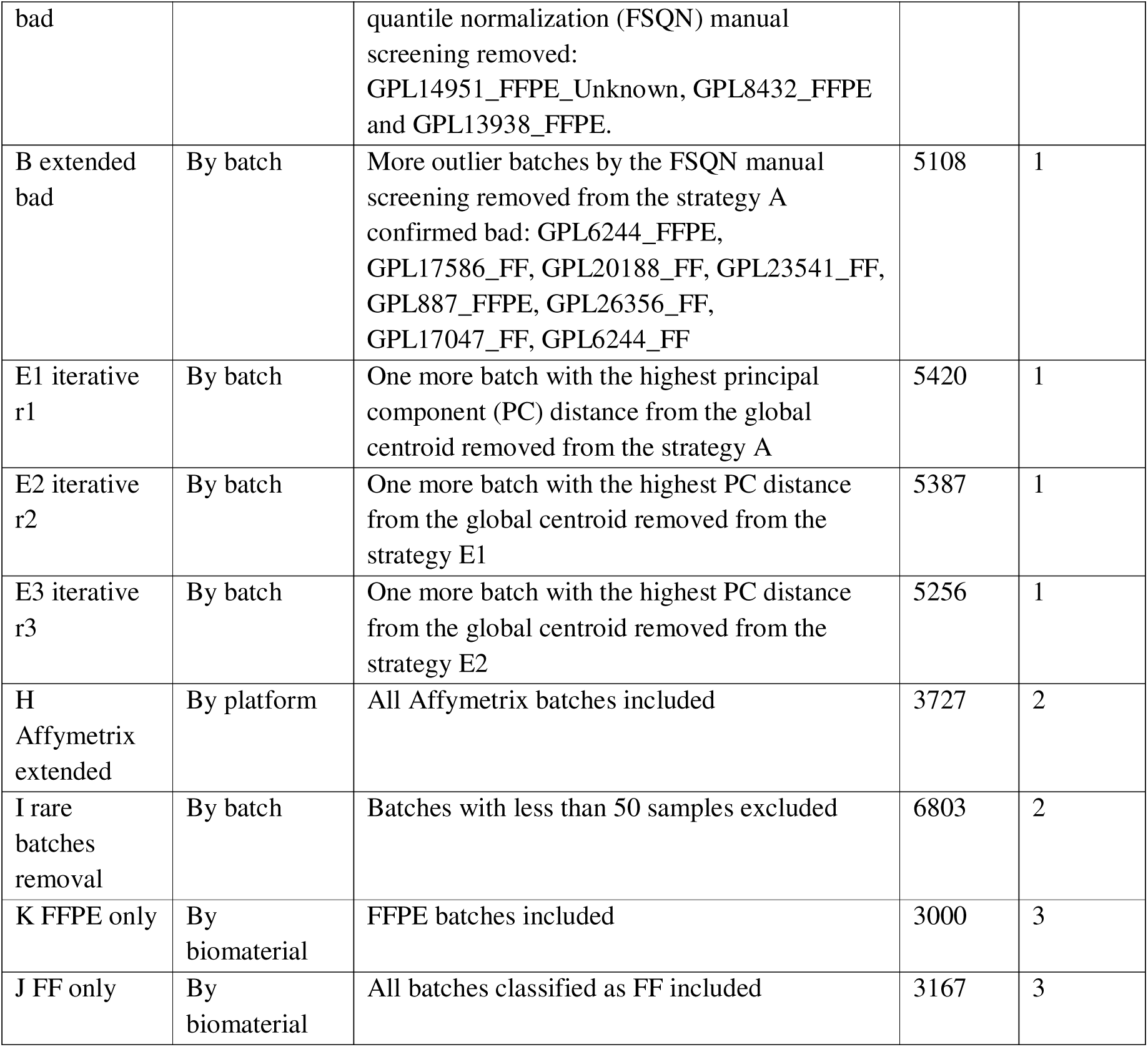
Prior batch removal strategies for the GC lymphoma dataset utilized in this study. The capital letters at the beginning of the strategy names are used as strategy labels throughout this paper.

The column ’Iteration when added’ in Table 1 records the pipeline development round in which each strategy was introduced. The initial ten strategies were evaluated in the first round; following results review, two additional strategies were added (Iteration 2). Analysis of the first two rounds established that the FF/FFPE biomaterial dichotomy was the most consequential factor in batch composition, motivating the addition of FF-only and FFPE-only strategies (Iteration 3).

### 2.6. Missing expression values imputation

Combining data from multiple platforms introduced extensive missing expression values because different microarray platforms interrogated different gene sets in the dataset (Supplementary File 2). We initially evaluated four imputation approaches: strict NA exclusion, k-nearest-neighbor (KNN) imputation, Softimpute, and missForest. The strict approach excluded all genes with at least one NA value across any sample in each strategy subset. missForest was excluded from the benchmark after exceeding 48 hours of computation time on a 32–48-core, 240 GiB RAM server without convergence, and was therefore deemed impractical for pipeline-scale application.

We ran KNN imputation using the KNNImputer from Python scikit-learn package version 1.3.2 (Pedregosa FABIANPEDREGOSA et al. 2011) with 5 nearest neighbors. For Softimpute we used the R library softImpute version 1.4-3 (Hastie et al. 2015) with default parameters.

Both KNN and Softimpute imputation were applied exclusively to genes for which fewer than 20% of samples had NA expression values, to ensure sufficient observed data for reliable imputation.

### 2.7. Harmonization methods

Through an in-depth literature search, we reviewed 75 publicly available harmonization methods and rejected 36 of them prior to implementation (Table 2).

**Table 2.** Rejection reasons for harmonization methods that were not taken forward to the implementation phase.

| Rejection reason | Methods number | Methods list |
| --- | --- | --- |
| Wrong data modality or architecture for the current dataset: scRNA-seq-only, deconvolution instead of expressions correction, cross-species, multi-omics-only, need raw counts/CEL/FASTQ files. | 13 | HARP, MoDAmix, scBatch, RNABC, CSN, fRMA, scGen, scVI, DESC, AutoClass, iNMF, MultiBaC, POIBM |
| Requires matched, paired or bridge samples not present in the data. | 7 | MatchMixeR, COCONUT, ESLR, QN-CN(CrossNorm), Ratio-based (Quartet), BRIDGE, Remeasure |
| Output is not a corrected expression matrix — it is an embedding, a probability, a category, or a score. | 5 | UPC, QD, NorDi, PLIDA, Divergence Analysis |
| Redundant with, inferior to or superseded by an adopted method. | 7 | Rank-in, QNR, Shambhala-1 (Borisov et al. 2019), QN-Z, RUV-2, BMC, Z-scaling (implemented as FSMVN) |
| No maintained or accessible software at screening time | 3 | IBN, DisTran, GQ |
| Preprocessing step, not a batch-correction method | 1 | Global Z-scoring |

After the rejection, we identified 39 harmonization methods potentially applicable to the current GC lymphoma dataset (Supplementary Table S1 in Supplementary File 2). We successfully implemented 33 of these methods in a Python 3.11 virtual environment, preferring native R package wrappers via rpy2 where R-specific implementations existed, and implementing standard methods (median scaling, quantile normalization) as custom Python functions. Except where Supplementary Table S1 states otherwise, we used the methods with their default parameters, to avoid further inflation of the ComboBatch pipeline combinatorial space. For SVA, ComBat, ComBat-seq, M-ComBat, InMoose ComBat-seq, ARSyN and Harman we supplied the annotation column ‘Diagnosis_cell_type_unified’ as the biology covariate in the model matrix, to protect its variance in the gene expression space. limma removeBatchEffect was run with the batch vector only and no design matrix, and RUV was run on ten housekeeping control genes with k = 2 and no biological covariate; neither implementation therefore protected the biology column. The parameters passed to every method are listed in Supplementary Table S1.

Six methods failed implementation and are documented with reasons in Supplementary Table S1. For one of the methods, Shambhala-2, (Borisov et al. 2022), we did extensive refactoring and containerizing, released it as a separate GitHub repository accessible via the link (https://github.com/Nikit357/Shambhala2_fast) and named 20_shambhala in the current analysis convention (Supplementary Table S1).

Each implemented method was assigned a qualitative harshness score (low, medium or high), from the published description of the algorithm and its source code. The criteria were applied in the following order. Low harshness: the method applies an additive or otherwise monotone transformation that leaves the rank order of genes within a sample unchanged (median scaling, limma, TMM, variance stabilization, SVA). Median scaling was the simplest among the low harshness methods: for each batch and each gene, its median was subtracted in the log_2_-transformed space. Medium: the method estimates a model or a small number of latent factors from the data and subtracts their contribution, or applies local neighbor-dependent offsets, so that the shape of the per-sample expression distribution is largely preserved (ComBat and its variants, RUV, MNN, Harmony, Scanorama, DWD, Harman). High: the method rewrites the per-gene or per-sample quantile function, replaces expression values by ranks, or changes the dimensionality of the data (quantile normalization, FSQN, rank normalization, TDM, NPN, AMDBNorm, exploBATCH, Shambhala-2). The tier assigned to each method and the reason for it are given in Supplementary Table S1, as well as linearity status of each method. Harmonization methods were incorporated across three development iterations; each successive iteration had a higher implementation failure rate consistent with their increasing algorithmic complexity (Supplementary Table S1). Among the methods tested, FSMVN and median scaling were the simplest, transforming biology to the least extent.

### 2.8. Post-removal

As an optional step after harmonization, we implemented a post-removal step on the premise that harmonization may fail for individual cohorts, which can then be excluded. For each of the batches in a dataset (by the RNA_BATCH cohort, maximum 29 batches) we calculated Euclidean distance of its centroid from the total harmonized dataset centroid in the principal component analysis (PCA) space (first 10 PCs), and we removed the batch with the highest centroid distance. This was done for each combination of batch removal strategy, imputation and harmonization method.

### 2.9. Harmonization pipeline architecture

We built the harmonization pipeline based on python scripts that can be found in the GitHub repository via the link https://github.com/Nikit357/FL_harmonization/tree/main/harmonization-scripts. The pipeline consisted of the two steps: 1) batch removal plus imputation (followed by log-transform) and 2) harmonization plus post-removal. Each step was orchestrated by a separate dispatcher script and executed by an individual job script, importing the necessary functions from the bench script. The modular implementation allowed additional combinations of the two steps to be run if necessary. Outputs of both steps were saved to Amazon Web Services (AWS) S3 storage as gzipped expression tables accompanied by annotation files.

### 2.10. Harmonization quality metrics

For harmonization quality assessment, we assembled a set of 87 polarity-defined scoring metrics together with 257 additional metrics computed but not included in the composite scoring (Supplementary Table S2 in Supplementary File 2, the full list of metric names in the sheet Metric_polarity in the same file). The additional metrics were needed as metadata, intermediate numbers for the final 87 scoring metrics calculation and as non-scoring measures of the harmonization output properties: number and fraction of zero genes and genes with expression below one, as well as gene expression percentiles (1^st^, 5^th^, 25^th^, 50^th^, 75^th^, 95^th^ and 99^th^), minimum, maximum and standard deviation. An additional set of biology classes prediction and biomarker expression preservation metrics (groups L, M, N) were used for the independent assessment of best approaches but were excluded from the composite scoring. The metrics were collected from the published literature on bulk, single cell and spatial transcriptomics batch correction (Supplementary Table S2), and their source code and seed numbers are available in GitHub via the link https://github.com/Nikit357/FL_harmonization/tree/main/harmonization-metrics-calculation. 633 biomarker genes for correlation were extracted from earlier FL and DLBCL studies (Dybkær et al. 2015; Kotlov et al. 2021; C. Sun et al. 2025), and a narrow set of 56 genes was selected to be present in all the batch removal strategies under the strict NA handling and having less than 20% of approaches with gene expression below 1 (Table_S3_gene_panel in Supplementary File 2).

The 87 scoring metrics spanned four major groups: (i) global distance metrics, quantifying the overlap of batch or biology classes in global PCA space; (ii) local neighborhood metrics, assessing the homogeneity of class intermixing in the neighborhood of each individual sample; (iii) distributional similarity metrics, evaluating the degree of convergence of per-cohort gene expression distributions after harmonization; and (iv) other metrics, including gene- and sample-level NA rate, fraction of bimodal cohorts and zero expression rate.

Harmonization quality metrics were computed and statistically analyzed using the scipy (v1.12.0) (Virtanen et al. 2020), scikit-learn (v1.3.2) (Pedregosa FABIANPEDREGOSA et al. 2011), statsmodels (v0.14.1) (Seabold and Perktold 2010), and umap-learn (v0.5.5) (McInnes et al. 2018) Python packages. 95% confidence intervals (CI) were assessed for Spearman correlations using the scipy.stats.bootstrap method, and for R^2^ values by harmonization parameter by T-statistic via scipy.stats.t.interval. All stochastic steps used a fixed random seed of 42. Principal component analysis was computed on standardized expression values; UMAP embeddings used n_neighbors = 30, min_dist = 0.3, two components and the Euclidean metric; tSNE embeddings used perplexity = 30, two components and the Barnes-Hut approximation, both computed on the principal component coordinates. The 1,000 genes sampled for the Kolmogorov-Smirnov metrics were drawn once with the same seed and reused for every approach.

### 2.11. Figures preparation

Basic plotting was done using the general Python packages seaborn (v0.13.2) (Waskom 2021) and matplotlib (v3.10.8) (Hunter 2007). Dimensionality reduction using PCA, UMAP and tSNE was done using scikit-learn (v1.3.2) (Pedregosa FABIANPEDREGOSA et al. 2011) and umap-learn (v0.5.5) (McInnes et al. 2018) Python packages. Individual panels were assembled into final figures in Figma (Figma: The collaborative canvas for design, code, and AI). Dendrograms for clustermaps were built in Euclidean space using the Ward linkage method (Strauss and Von Maltitz 2017).

### 2.12. Basic data analysis software

Data analysis and table manipulations were done using pandas (v2.3.3) (Mckinney 2010; team) and numpy (v1.26.4) (Harris et al. 2020).

### 2.13. AI usage

Claude Code with the Claude Sonnet 4.8, Sonnet 5 and Opus 5 models (Liu et al. 2026 Apr 14) was used for preliminary scientific literature review, code writing and refinement and draft text grammar and language checks. All literature review and code writing was done in a plan-then-act mode: the AI agent first wrote a research and implementation plan, the authors then reviewed and commented on it, and after two to five review cycles the agent implemented the requested feature. AI specifications can be found in the project repository (https://github.com/Nikit357/FL_harmonization) as CLAUDE.md files in all the subfolders. The intermediate research outputs and implementation plans, as well as additional documentation are stored in other markdown files that contain the following key words in their names: ‘project_overview’, ‘plan’, ‘implementation’, ‘research’. The Claude code skills generated during the article preparation are stored in the repository root directory in the path ‘.claude/skills/’ as markdown files with human readable instructions.

Iterative code refinement for complicated visuals was done using Google Gemini 3.1 Pro (Omar et al. 2025). The research study purpose, objectives and design, the ComboBatch architecture, text writing and figures assembly were done manually by authors.

## 3. RESULTS

### 3.1. Application of batch removal strategies, harmonization methods, imputation methods and post-removal to the FL dataset

We assembled a multi-platform dataset of 7,174 bulk transcriptomic profiles from 88 cohorts spanning RNA-seq, Affymetrix, Illumina, and Agilent microarray platforms (Figure 1A). With a focus on GC B-cell malignancies, the dataset comprised 4,466 DLBCL, 1,697 FL, and 889 normal B-cell samples (Figure 1B), together with 88 Burkitt lymphoma and 34 high-grade B-cell lymphoma samples from overlapping cohorts serving as additional reference biology groups. Complete sample annotation is provided in Supplementary File 1.

**Figure 1.**
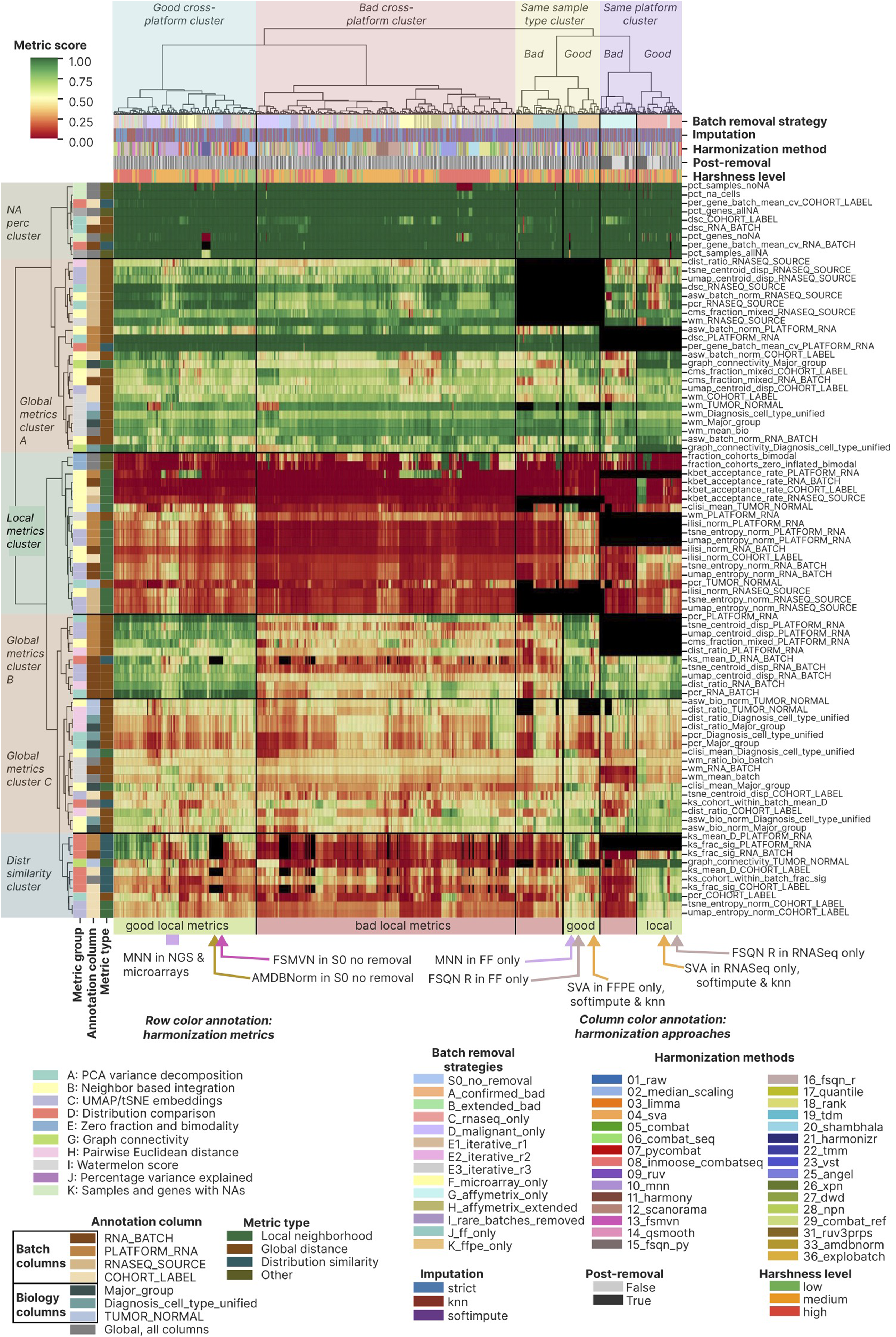
Dataset composition and computational pipeline. **(A)** Dataset composition by batches. **(B)** Dataset composition by biology groups, each point represents one sample here and in the panel A. **(C)** Scheme of the ComboBatch computational pipeline. **(D)** Bar plot of sample counts by batches and biology, ordered by batches. Biology groups are color coded according to the B panel color legend. **(E)** Bar plot of sample counts by biology and batches, ordered by biology groups. Batches are color coded according to the A panel color legend. **Alt text:** Multi-panel overview of the dataset and pipeline: bubble plots of sample composition by batch and by biological group, a schematic of the nine-step ComboBatch pipeline, and stacked bar plots of sample counts by batch and by biology.

To benchmark publicly available harmonization tools in an unbiased manner and identify the optimal approach for a dataset with subtle biological and pronounced technical differences, we designed a nine-step computational pipeline named ComboBatch (Figure 1C). Starting from the full multi-platform dataset, ComboBatch applied one of 14 prior batch removal strategies — retaining sample subsets defined by platform or biomaterial type or excluding outlier batches identified by annotation or unsupervised screening — followed by imputation of missing expression values using KNN or Softimpute (or strict exclusion of genes with any NA), log2-transformation per cohort, and one of 33 harmonization methods (Supplementary Table S1). Optionally, the pipeline excluded a single PCA-outlier batch after harmonization as a fine-tuning step. The pipeline then computed 87 polarity-defined scoring metrics and assembled an integrated clustermap across all harmonization approaches — each defined as a unique combination of prior batch removal strategy, imputation method, harmonization algorithm, and post-removal status. The full cross-product of these four factors yielded 2,234 successfully completed harmonization approaches that had fewer than 5% of samples with entirely missing gene expression values (by the condition pct_samples_allNA < 5), out of 2,407 approaches with harmonization completed with any resulting gene expression matrix and out of 2,772 theoretically possible approaches. The methods 34_arsyn and 38_harman had all the harmonization approaches with more than 5% samples with NA only expression, so we excluded them from the downstream analysis and studied the final set of 31 methods. The metrics by all approaches can be accessed in Supplementary File 3.

The dataset was markedly imbalanced with respect to both batch and biology group composition. Among the six largest RNA batches by sample count, the fifth (GPL14951_FFPE, 810 samples) and sixth (GPL570_FFPE) contained DLBCL exclusively, while the second largest (GPL570_Unknown, 1,007 samples) comprised only four biological groups: FL, DLBCL, Burkitt lymphoma, and normal B cells (Figure 1D). The distribution of biological groups across batches was more uniform (Figure 1E): DLBCL and FL were represented in 12 and 15 batches, respectively, whereas rare groups showed extreme batch skew — high-grade B-cell lymphoma was confined to a single batch (RNASeq_FFPE_Exome_capture), and plasmablasts and Burkitt lymphoma were each represented by two batches. Analogous composition skew was observed at the cohort level (Supplementary Figure 1): the five largest cohorts by sample count were exclusively DLBCL (Supplementary Figure 1A), except for the SOM cohort (Loeffler-Wirth et al. 2022). Normal B cells were profiled predominantly in smaller cohorts. With two exceptions (Kassandra and PUB_FL_NCISTAUDT), each cohort contributed samples to a single RNA batch (Supplementary Figure 1B).

The initial step of ComboBatch was prior batch removal, implemented across 14 distinct strategies (Figure 2A), grouped by selection criterion: platform type (RNA-seq only; Affymetrix-only microarrays, all microarrays), biomaterial (FF-only; FFPE-only), tumor biology (malignant only), and outlier batch exclusion (rare batches and those performing poorly in initial FSQN screening; Supplementary Figure 1C). Imputation substantially increased the number of genes available for downstream analysis (Figure 2B): strict NA exclusion yielded between 3,447 genes (S0 and D strategies) and 6,797 genes (C strategy), whereas KNN and Softimpute imputation extended this range up to 15,885 (C strategy) genes. KNN and Softimpute produced identical gene sets because both were applied exclusively to genes with fewer than 20% NA values per strategy subset.

**Figure 2.**
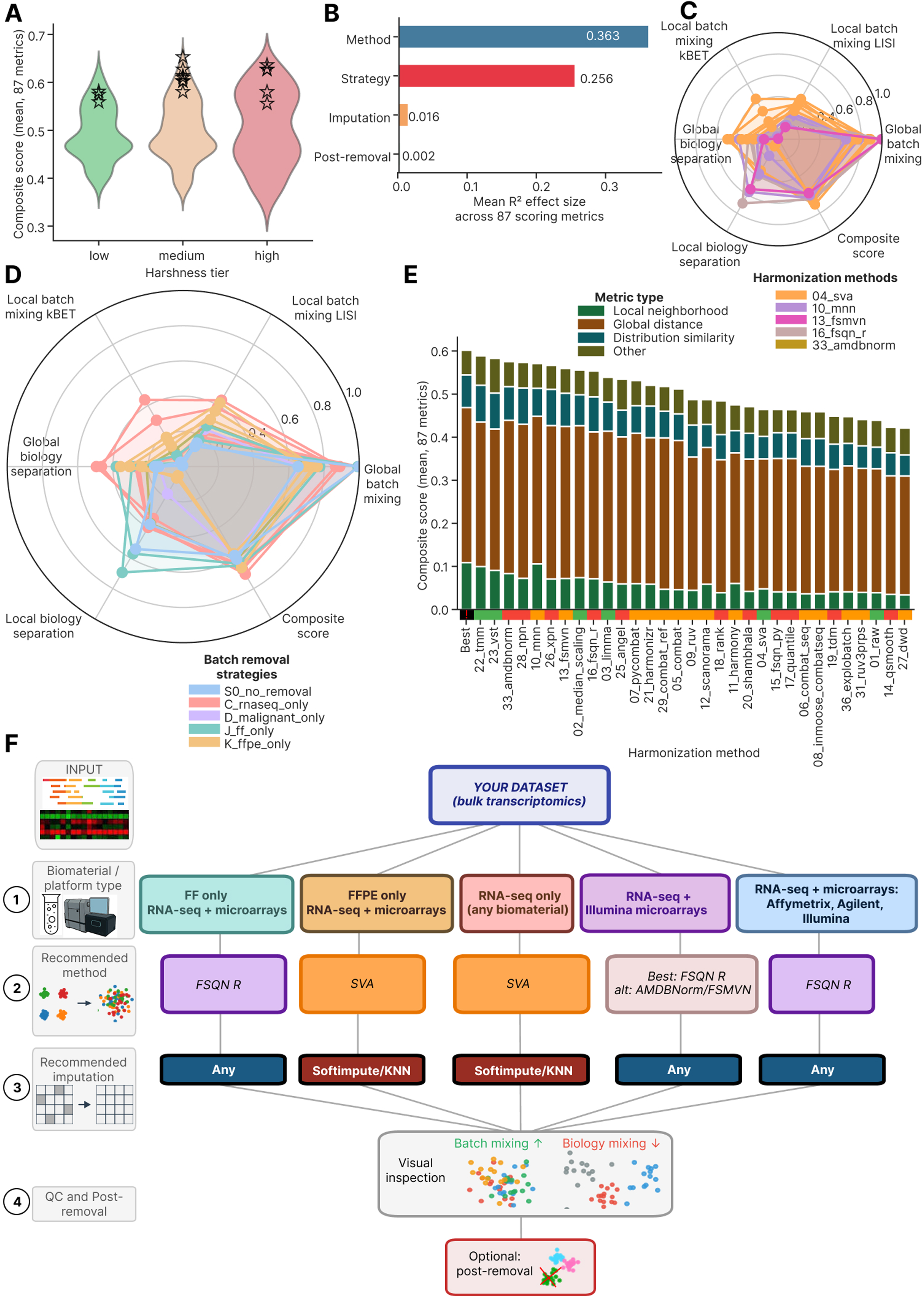
Batch removal strategies, NA genes and imputation. **(A)** Circos-Sankey plot showing batch removal strategies and number of samples remaining after each strategy being applied. The strategies extend from the central blue circle symbolizing the S0 strategy, and logically they are divided into the four groups (arcs in the blue circle): by platform, by biology, by biomaterial, bad batches removal. **(B)** Bar plots showing gene number under the strict approach (bottom plot, all genes with at least one NA in a dataset removed) and after the imputation procedures (top plot, KNN or Softimpute). Gene number counted as number of genes having non-NA expressions in at least one sample in the dataset. **(C)** Heat map of NA/defined expression by genes and samples in the dataset. Samples and genes are ordered by ascending NA count from left to right and from top to bottom. Palettes for color annotation in the top of the heat map can be found in Figure 1A, 1B. **(D)** Circos-Sankey plot showing gene numbers and overlapping of gene sets for batch removal strategies under the strict NA handling and gene imputation. Batch removal strategies are structured by the same four groups as in the panel A. **Alt text:** Four panels on batch-removal strategies and missing data: a Circos–Sankey plot of the 14 strategies with remaining sample counts, bar plots of gene numbers before and after imputation, a heat map of defined versus missing expression across genes and samples, and a Circos–Sankey plot of gene-set overlap between strict and imputed strategies.

Strict NA exclusion produced predominantly bimodal (72 of 88; Supplementary Figure 2) and unimodal expression distributions (16 of 88), the former tending to associate with RNA-seq cohorts and the latter being connected with the microarray’s ones (Supplementary Figure 2). KNN and Softimpute yielded more homogeneous per-cohort distributions, reducing the number of cohorts with unimodal distributions to 5 (KNN, Supplementary Figure 3) and 3 (Softimpute, Supplementary Figure 4), respectively. Notably, Softimpute also introduced spurious trimodal expression distributions in 14 cohorts (Supplementary Figure 4), an imputation artifact consistent with the low-rank approximation underlying the algorithm and its sensitivity to heterogeneous missingness patterns (Hastie et al. 2015).

The initial dataset exhibited substantial heterogeneity in gene coverage before any NA handling (Figure 2C): RNA-seq, GPL570, and GPL14951 batches provided expression values for more than 15,000 genes, whereas rare platform batches (GPL20188_FF, GPL13158_FF, and GPL16686_FF) contained only 6,000-7,000 genes with defined expression. This inter-platform heterogeneity was the primary driver of the restricted gene numbers obtained under strict NA exclusion (3,447-6,797 genes), a limitation largely overcome by imputation (6,797-15,885 genes).

We visualized gene set overlaps across imputation and strict NA-exclusion approaches using a Circos-Sankey plot (Figure 2D). Iterative batch removal — in PCA space (strategies A to E1 to E2 to E3) or restricted to microarrays (strategies F to H to G) — did not substantially increase post-imputation gene counts, although strict gene numbers rose from 3,447 (strategy F) to 6,133 (strategy H), and from 3,520 (A) to 5,658 (B) and 5,175 (E1). By contrast, imputation yielded a 3.4-fold gene count increase for the unrestricted S0 strategy (3,447 to 11,768 genes). The strategies with the highest post-imputation gene numbers (A, E1-E3, B, D, K, C) shared near-identical gene sets, with pairwise Jaccard indices of 95-100% (Supplementary Figure 5), indicating that imputation is most effective when the contributing platforms and biomaterials have broadly overlapping transcriptome coverage.

### 3.2. Best approaches selection

For each of the 2,234 successful harmonization approaches, we computed up to 344 harmonization metrics per approach, selected from the published literature to span all dimensions of cross-platform harmonization quality (Supplementary File 3; Supplementary Table S2). For strategies restricting samples to a single platform or biomaterial (strategies C, G, J, and K), a subset of annotation-dependent metrics was not defined: for example, in strategy J (FF only), the column RNASEQ_SOURCE was invariant, rendering all metrics conditioned on that column inapplicable. This resulted in decreased non-NA metrics for strategies C, G, J, and K (Supplementary Figure 6). Additionally, some approaches were terminated by a 3-hour CPU-time limit imposed by the job scheduler – these approaches are marked as red in Supplementary Figure 6.

For 87 of the 344 computed metrics, we manually assigned polarity values of +1 (higher is better) or - 1 (lower is better), based on the principle that high-quality harmonization should maximize batch mixing and preserve biological group structure: metrics quantifying the homogeneity of batch-class distribution in expression space were assigned positive polarity, whereas metrics quantifying biological group mixing were assigned negative polarity. For the remaining 257 metrics polarity was set as 0 as they did not participate in the final scoring. Polarity assignments for all the 344 metrics are listed in Supplementary File 2, sheet ‘Metric_polarity’; the metric names there match the column names of Supplementary File 3.

Using the 87 polarity-defined metrics and 2,234 qualifying harmonization approaches, we constructed a clustermap to characterize the overall quality landscape of the harmonization approach space (Figure 3). At the level of metric structure, the 87 metrics resolved into six co-clustering groups: three comprising global metrics, and three comprising distributional similarity, NA-percentage, and local mixing metrics, respectively. To characterize the relationship between local and global metric performance, we constructed a separate Spearman cross-correlation clustermap among all 87 metric vectors computed across the 2,234 approaches (Supplementary Figure 7). This analysis revealed an isolated cluster of 33 metrics — 22 of which were local mixing metrics — that negatively correlated with a second cluster of global metrics (10 global, 6 distributional, and 1 other). This could point at a fundamental trade-off between the global batch separation and local neighborhood batch mixing. A third cluster showed no dominant correlation direction with the other two and consisted of global, other and distributional metrics. Within the larger metric-type clusters, smaller sub-clusters formed around shared annotation columns (primarily PLATFORM_RNA and RNASEQ_SOURCE), with no systematic grouping by metric group. PCA, UMAP, and tSNE embeddings of metric vectors (Supplementary Figure 8) confirmed large-scale co-localization by metric type and finer-scale grouping by annotation column, with no pattern by metric group.

**Figure 3.**
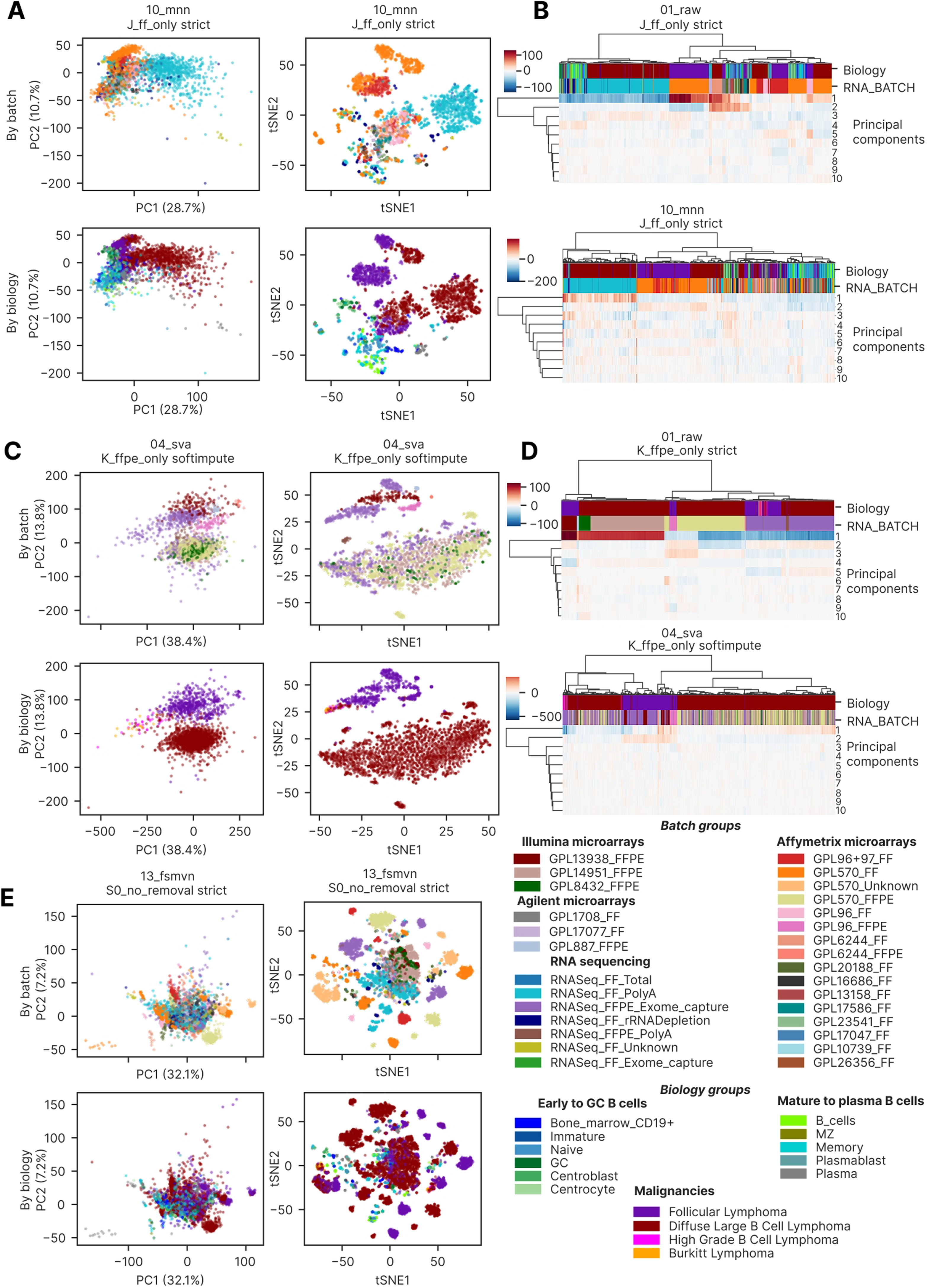
Clustermap of 2,234 harmonization approaches (rows) in a space of 87 quality metrics (columns), polarity adjusted and min-max scaled. Black cells represent NA metrics. The best approaches identified from the clustermap are indicated beneath it by arrows, with a rectangle marking the MNN group. **Alt text:** Clustermap of 2,234 harmonization approaches (columns) scored on 87 quality metrics (rows); black cells mark missing metrics, and the best-performing approaches are indicated beneath the map.

At the level of harmonization approaches, the clustermap revealed four major clusters (Figure 3). Based on relative metric performance and the co-clustering patterns of batch removal strategies, we designated these clusters: ’good cross-platform’, ’bad cross-platform’, ’same sample type’, and ’same platform’. The ’same sample type’ and ’same platform’ clusters each contained a good and a bad sub-cluster, separated primarily by local mixing metrics — specifically LISI values and UMAP/tSNE entropy. The ’good cross-platform’ cluster correspondingly showed higher local metric values than its ’bad’ counterpart, with two-sided Mann-Whitney FDR-corrected p < 0.05 for 20 / 22 local metrics investigated between the two clusters and log_2_(fold change of the medians in two clusters) being in the interval 0.26 – 3.7. Across all harmonization approaches, kBET acceptance rate was the only local mixing metric that remained near-zero universally (RNA_BATCH median 0.0022 in ’good cross-platform’ cluster and 0 in the ‘bad’ one), with the sole exception of RNA-seq-only configurations with post-removal applied.

Metrics in the ’NA percentage’ cluster showed uniformly high performance across all approaches; the sole exception was pct_genes_noNA, which recorded poor performance exclusively for the 21_harmonizr harmonizer: 0% for the strategies A, B, C, E1-E3, J and S0, and 87.2–100% for the remaining strategies. Metrics conditioned on the RNASEQ_SOURCE and PLATFORM_RNA annotation columns were undefined for strategies restricted to a single sample type (J, K) or a single platform (G, C), respectively, because these columns were invariant within those strategy subsets. Global cluster B metrics (primarily tSNE/UMAP centroid-based measures) showed highest performance within the ’good cross-platform’ cluster, the good sub-cluster of the ‘same sample type’ cluster and the entire ‘same platform’ cluster: two-sided Mann-Whitney FDR-corrected p < 0.05 for 9 / 10 metrics between the ‘good’ and the ‘bad’ cross-platform cluster, log_2_(fold change of medians) being in the interval 0.51 – 2.0. Distributional similarity metrics showed an analogous pattern, while global cluster C metrics showed intermediate performance across the full approaches set.

We calculated Spearman cross-correlations between all harmonization approaches in the 87-metric scoring space and constructed a clustermap of these correlations (Supplementary Figure 9). The four clusters identified in the primary clustermap — ’same platform’, ’same sample type’, ’good cross-platform’, and ’bad cross-platform’ — were reproduced, confirming that cluster identity was not an artifact of the primary metric-space embedding. As in the primary clustermap, clustering was driven primarily by batch removal strategy and secondarily by harmonization method. Unlike the metric cross-correlation clustermap (Supplementary Figure 7), no negatively correlating approach clusters were observed. The ’same sample type’ cluster showed the strongest deviation from all other ones, with median Spearman correlation of 0.554 (median absolute deviation (MAD) 0.104) relative to ’same platform’ and ’good cross-platform’ clusters, whereas the remaining correlations had a median of 0.730, MAD 0.129 (two-sided Mann-Whitney p-value < 10^-200^ between the two groups, log_2_ fold change of the medians 0.40). PCA, UMAP, and tSNE embeddings of harmonization approaches in the 87-metric space (Supplementary Figure 10) confirmed the dominant influence of batch removal strategy, the lower importance of harmonization method and revealed no co-clustering by method harshness.

### 3.3. Best approaches

Based on the clustermap visual inspection combined with quantitative composite scoring, we identified eight best-performing harmonization approach groups (15 individual approaches, Table 3), spanning multi-platform and single-platform strategies as well as FF-only and FFPE-only compositions. By the clustermap metrics, MNN and SVA emerged as the most versatile harmonization methods: MNN achieved top performance across multi-platform and FF-only strategies, while SVA showed highest performance in RNA-seq-only and FFPE-only contexts. FSMVN was the simplest method among the best approaches, since it was implemented as Z-scaling applied individually to each gene in each batch.

**Table 3.** The 15 best harmonization approaches (8 groups) identified via the integrative clustermap assessment in **Figure 3**.

| Group name | Batch removal strategies | Harmonization methods | Imputation | Reason to include | Approach names in the group (15 best approaches in total) |
| --- | --- | --- | --- | --- | --- |
| MNN multiplatform | Multiplatform strategies: S0, D, H | MNN | Any | Superior local metrics performance compared to the rest approaches in the ‘good multiplatform cluster’ | D_malignant_only_strict_10_mnn, H_affymetrix_extended_strict_10_mnn, S0_no_removal_strict_10_mnn |
| FSMVN in S0 | S0 (RNA-seq + Illumina microarrays scenario) | FSMVN | Any | Best performance by clisi_mean_Major_group in the Good cross platform cluster, mixing of NGS and Illumina microarrays on UMAP and tSNE | S0_no_removal_strict_13_fsmvn |
| AMDBNorm in S0 | S0 (RNA-seq + Illumina microarrays scenario) | AMDBNorm | Any | Best performance by clisi_mean_Major_group in the Good cross platform cluster, mixing of NGS and Illumina microarrays on UMAP and tSNE | S0_no_removal_strict_33_amdbnorm |
| MNN FF only | J | MNN | Any | Best performance by local metrics in the FF part of the good “Same sample | J_ff_only_strict_10_mnn |

|  |  |  |  | type” cluster |  |
| --- | --- | --- | --- | --- | --- |
| FSQN R FF only | J | FSQN R | Any | Near best performance by graph_connectivity_Diagnosis_cell_type_unified and clisi_mean_TUMOR_NO RMAL in the FF part of the good “Same sample type” cluster. Mixing of NGS and microarrays on UMAP and tSNE plots | J_ff_only_strict_16_fsqn_r |
| SVA FFPE only | K | SVA | Any but KNN and Softimpute are preferred | Best performance by local metrics in the FFPE part of the good “Same sample type” cluster | K_ffpe_only_knn_04_sva, K_ffpe_only_softimpute_04_sva, K_ffpe_only_strict_04_sva |
| SVA RNA-seq only | C | SVA | KNN, Softimpute | Best performance by local metrics in the RNA-seq part of the good “Same platform” cluster | C_rnaseq_only_knn_04_sva, C_rnaseq_only_softimpute_04_sva |
| FSQN R RNA-seq only | C | FSQN R | Any | High-quality mixing of RNA-seq FF and FFPE batches on UMAP and tSNE | C_rnaseq_only_knn_16_fsqn_r, C_rnaseq_only_softimpute_16_fsqn_r, C_rnaseq_only_strict_16_fsqn_r |

Without harmonization, batch removal and under strict NA handling, the FL dataset exhibited a pronounced batch effect structure in both PCA and tSNE space with 81.1% of the total variance explained by the first PC (Figure 4A), attributable mainly to batch effect. In PCA, RNA-seq samples formed a compact central cluster, while Affymetrix GPL570 batches arranged in a linear upper-right trajectory, and the remaining microarray batches (Affymetrix, Illumina, and Agilent) formed a lower-right trajectory. In tSNE, RNA batches dispersed into discrete compact clusters across the full embedding, confirming the validity of the RNA_BATCH annotation, with no discernible biological grouping. After MNN harmonization under strict NA handling, a single cohort outlier remained apparent in PCA space — the SOM cohort, which had undergone prior quantile normalization — while the remaining cohorts clustered together (Figure 4B). tSNE revealed only three batch outliers: a subset of RNASeq_FFPE_Exome_capture batch, a subset of GPL570_Unknown batch (the same SOM cohort), and the majority of the GPL570_FFPE batch. All remaining samples organized into biologically coherent groups, with FL, DLBCL, and normal B cells forming discrete clusters with batches mixing within each group, although Illumina microarray samples formed a distinct sub-cluster. Strategy D (malignant-only, Figure 4C) produced a further improvement in the concordance of biological clustering and batch mixing. Hierarchical clustering in the first-ten-PC space before and after MNN harmonization — with the SOM cohort removed — confirmed increased batch intermixing after harmonization: FL and normal B cells resolved into single isolated clusters, though DLBCL samples retained a partially batch-structured dendrogram topology (Figure 4D). Strategy H (Affymetrix extended) under MNN similarly highlighted the pre-normalized SOM cohort as a PCA outlier (Supplementary Figure 11A), and all three MNN-harmonized strategies showed 2-3 consistent batch outliers in both UMAP and PCA space (Supplementary Figure 11B, C).

**Figure 4.**
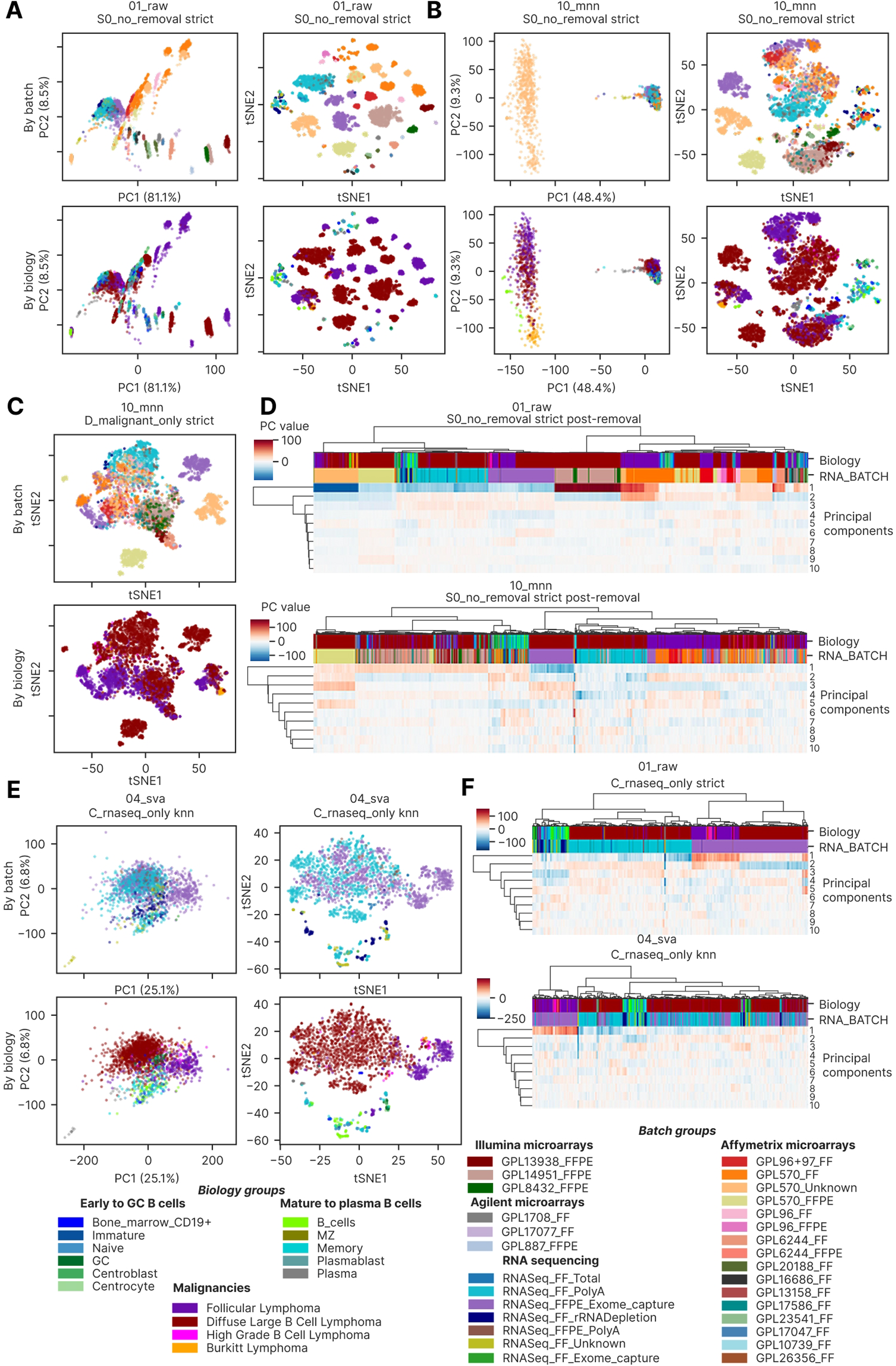
SVA in RNA-seq only and MNN in multiplatform strategy, the best harmonization approaches. **(A)** Raw dataset without batch removal (S0 strategy) dimensional reduction visualization in a 2 X 2 square grid. The upper row is visualization by batch, the lower one shows biological groups. The left column contains PCA plots, the right column shows tSNE plots. **(B)** PCA and tSNE visualization of batches and biology in the MNN harmonized S0 dataset, the structure is the same as in the panel A. **(C)** tSNE visualizations of MNN harmonized D dataset (malignant only), the lower one is by batch, and the upper one is by biology. **(D)** Comparison of first 10 PCs dendrograms in the raw (top) and MNN-harmonized S0 dataset (bottom), with color annotation by batch and biology. **(E)** Square grid with PCA/tSNE visualizations of SVA-harmonized KNN-imputed C dataset (RNA-seq only), the structure is the same as in the panel A. **(F)** Comparison of first 10 PCs dendrograms in the raw (top) and SVA-harmonized, KNN imputed S0 dataset (bottom), with color annotation by batch and biology. Color palettes of batch and biology groups are applicable to all the panels of this figure and are shown below the E and F panels. **Alt text:** Dimensionality-reduction and dendrogram panels showing that MNN (multi-platform) and SVA (RNA-seq only) harmonization mix batches while preserving separation of FL, DLBCL and normal B cells, relative to the unharmonized data.

We next examined the second top-performing approach: SVA applied to the RNA-seq only strategy (Figure 4E). Both PCA and tSNE revealed near-complete batch mixing — including the previously outlying RNASeq_FFPE_Exome_capture batch — with concurrent preservation of biological group separation. This level of performance was achieved exclusively with KNN and Softimpute imputation; strict NA handling failed to mix the major FF and FFPE batches in PCA, UMAP, or tSNE space (Supplementary Figure 12A-C, respectively), and the strict SVA result was visually indistinguishable from the unharmonized data. Hierarchical clustering in the first-ten-PC space before and after SVA harmonization (Figure 4F) demonstrated a near-complete transition from a batch-structured to a biology-structured dendrogram: FL samples resolved into a single discrete cluster, and normal B cells formed a sub-cluster nested within the broader DLBCL clade.

The second-best approach for the RNA-seq-only strategy (C) was FSQN R (16_fsqn_r) — the method implemented using the original R package, as distinct from the custom Python re-implementation (method 15_fsqn_py). On PCA, samples clustered by biological group (Supplementary Figure 13A), while non-linear dimensionality reduction revealed additional substructure within biological groups with improved batch mixing relative to the unharmonized data (Supplementary Figure 13B, C): there were three batch-derived subclusters in the DLBCL cluster and two in the FL cluster.

We next visualized and assessed the remaining top-performing approaches across all benchmark strategies (Figure 5). In the FF-only MNN-harmonized dataset, PCA revealed incomplete batch mixing but a biologically meaningful ordering of samples: FL between normal cells and DLBCL (Figure 5A). tSNE confirmed this dominance of biology over batch, but same-batch connectivity remained high: most NGS and Affymetrix batches formed isolated clusters (Figure 5A). Yet, clustering in the first 10 PCs revealed an improvement of batch mixing primarily for microarrays (Figure 5B). FSQN R harmonization was selected as the second-best approach for FF only, because biological structure dominated less clearly in PCA, UMAP and tSNE (Supplementary Figure 14A, B, C, respectively). FL batch-level subclusters surrounded the DLBCL cluster instead of being between it and normal cells (Supplementary Figure 14A).

**Figure 5.**
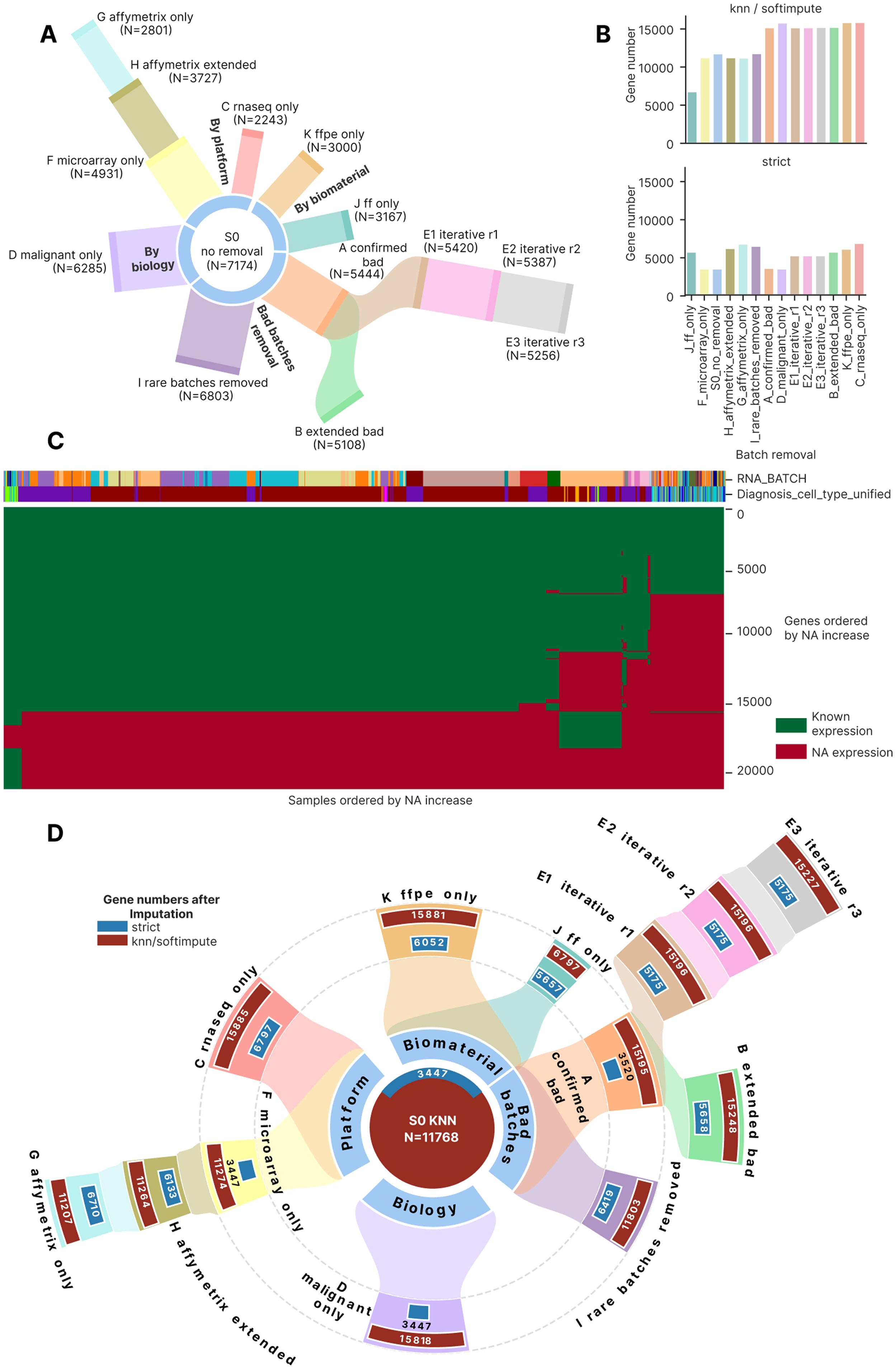
The second part of the best harmonization approaches: MNN for FF only (J), SVA, Softimpute for FFPE only (K), FSMVN for S0 with Illumina microarrays and NGS. **(A)** FF only batch removal harmonized with MNN dimensional reduction visualization in a 2 X 2 square grid. The upper row is visualization by batch, the lower one shows biological groups. The left column contains PCA plots, the right column shows tSNE plots. **(B)** Comparison of first 10 PCs dendrograms in the raw (top) and MNN-harmonized FF only dataset (bottom), with color annotation by batch and biology. **(C)** PCA and tSNE visualization of batches and biology in the SVA harmonized Softimpute FFPE only dataset, the structure is the same as in the panel A. **(D)** Comparison of first 10 PCs dendrograms in the raw (top) and SVA-harmonized, Softimpute FFPE only dataset (bottom), with color annotation by batch and biology. **(E)** PCA and tSNE visualization of batches and biology in the FSMVN harmonized S0 dataset, the structure is the same as in the panel A. **Alt text:** PCA, tSNE and dendrogram panels for further top approaches: MNN for fresh-frozen only, SVA with Softimpute for FFPE only, and FSMVN for RNA-seq combined with Illumina microarrays.

For the FFPE only batch removal strategy, the top approach of SVA harmonization and Softimpute imputation was more successful: DLBCL samples from NGS, Affymetrix and Illumina microarray batches were fully intermixed in both PCA and tSNE (Figure 5C), whereas FL batch-dependent subclusters formed a large cluster without intersection with DLBCL. Clustering on the first 10 PCs (Figure 5D) produced two high-level branches, each consisting of FL and DLBCL and having homogeneous mixing of batches in DLBCL and heterogeneous in FL. Before harmonization we observed a complete dominance of batch over biology on the dendrogram (Figure 5D). Softimpute imputation mixed batches in FFPE only better than the KNN and strict approach, leading to the minor Illumina microarray batch GPL8432_FFPE mixing with other batches in Softimpute. This effect was not clear on PCA (Supplementary Figure 15A) but was observable on UMAP and tSNE (Supplementary Figure 15B, C, respectively).

Finally, FSMVN and AMDBNorm methods showed promising results for the cross-platform harmonization of Illumina microarrays and NGS. FSMVN in S0 led to co-clustering of the large batch RNASeq_FF_PolyA and Illumina microarrays on tSNE (Figure 5E). As previously, the effect was not observed on PCA plots for any imputation (Supplementary Figure 16A) but was observable on UMAP and tSNE for all imputations (Supplementary Figure 16B, C, respectively). AMDBNorm mixed the RNASeq_FF_PolyA and Illumina microarrays batches on PCA, UMAP and tSNE (Supplementary Figure 17A, B, C, respectively). The effect was expressed to the highest extent in the S0 strategy, whereas FF and FFPE only strategies led to moderate success (Supplementary Figure 17).

### 3.4. Relative impact of batch removal strategies, harmonization methods, imputation methods and post-removal

To integrate the multi-dimensional metric signals, we computed a composite harmonization score as the mean of polarity-normalized values across all the 87 scoring metrics, weighted by their polarity (Figure 6A). We compared the composite score between harmonization methods by their harshness: the high harshness methods had higher maximum values (0.69 compared to 0.66 for low harshness methods) but had comparable medians: 0.479 for low harshness approaches, 0.488 for medium and 0.493 for the high ones (MAD 0.044 in the whole dataset). Despite the close medians, low harshness approaches significantly differed from both the medium and the high ones (two-sided Mann-Whitney FDR-corrected p = 0.041 for both comparisons). The clustermap best methods occupied the upper 1/3 of the distributions, having the composite score 0.559 and higher (median 0.603, MAD 0.023).

**Figure 6.**
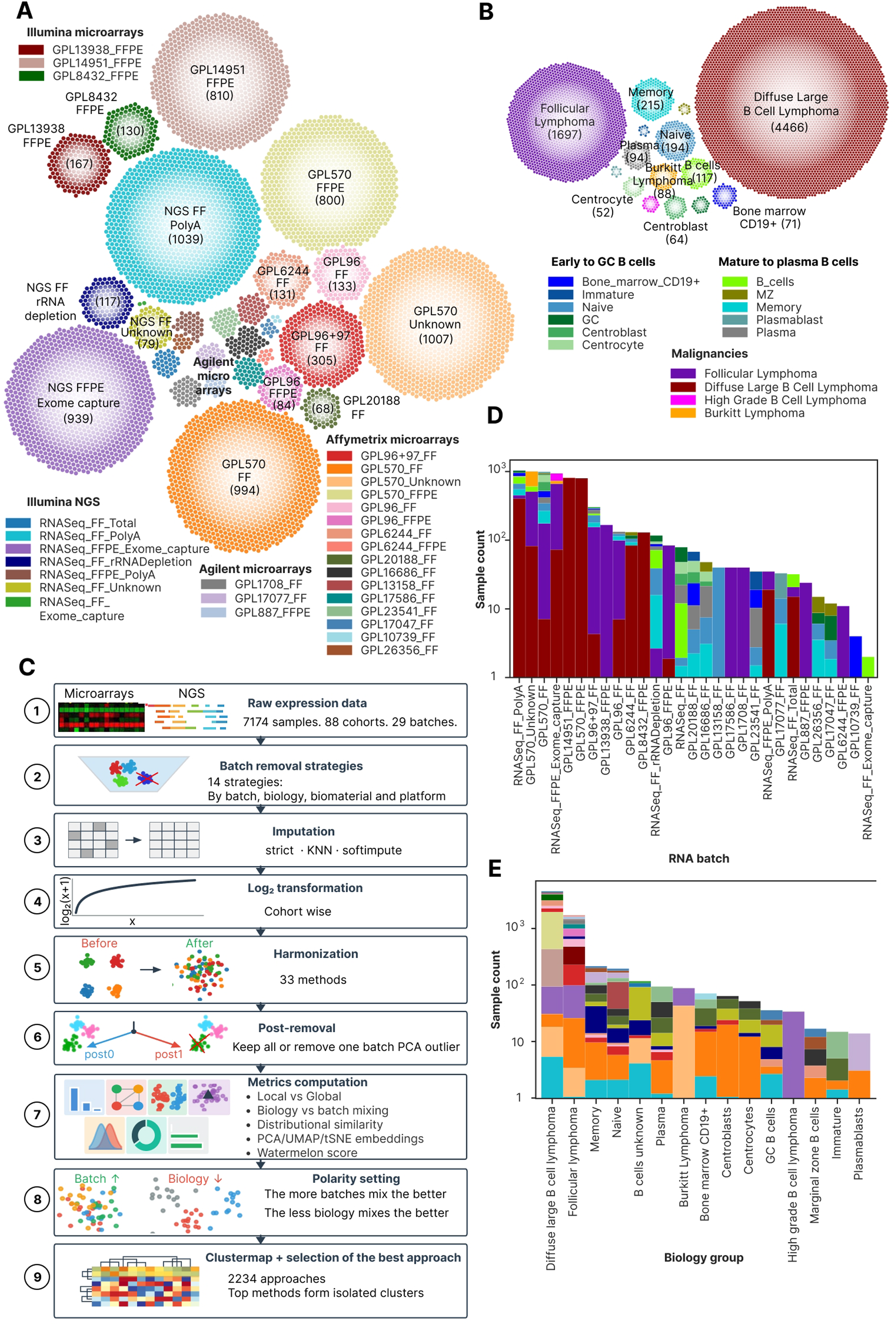
Composite performance score, clustermap best approaches behavior and the final decision tree. (A) Violin plot showing the composite score by harmonization method harshness level, with the clustermap best approaches shown in black stars. (B) Horizontal barplot showing R² effect sizes of the composite score for four experimental factors: harmonization method, batch removal strategy, imputation method, and post-removal. (C) Radar plot showing performance of the clustermap best approaches by six axes (mean normalized metrics of each group): local batch mixing by kBET, local batch mixing by LISI, local biology preservation, global biology preservation, global batch overlapping and the total composite score. The approaches are colored by their harmonization methods. (D) Enlarged radar plot showing the same best approaches with the same axes as in panel C but colored by batch removal strategy. (E) Stacked bar chart of cumulative normalized score per harmonization method, decomposed by metric type (local neighborhood, global distance, distributional similarity, other). Y-axis represents cumulative summed normalized score across all metrics of each type. (F) Data-driven decision tree showing the workflow and the best harmonization approaches depending on the dataset composition. At the biomaterial/platform type (step 1) we consider only the cross-platform and/or cross-biomaterial datasets. At the imputation step (step 3), ‘Any’ means both imputation methods (KNN and Softimpute) and no imputation (the strict approach). **Alt text:** Composite-performance figure: violin plot of composite score by method harshness, bar plot of R-squared effect sizes for the four factors, radar plots of the best approaches across six performance axes, a stacked bar chart of score by metric type, and the final harmonization decision tree.

To quantify the relative contribution of each experimental factor to overall harmonization quality, we computed the R² effect size of each factor separately for each of the 87 scoring metrics across the 2,234 successful approaches and averaged it over the metrics (Figure 6B). Harmonization method emerged as the dominant factor, explaining on average 36.3% of the variance of a single quality metric (95% CI 31.5, 41.1), followed by batch removal strategy (R² = 0.256, 95% CI 0.208, 0.305). Imputation method contributed only 1.6% of variance (95% CI 0.9, 2.2%), and post-removal contributed only 0.2% (95% CI 0.1, 0.3%), confirming the four-factor hierarchy: method > strategy > imputation > post-removal. It should be noted that earlier in the main clustermap (Figure 3) we visually observed the priority of batch removal strategy, followed by harmonization method, but post-removal and imputation comprised a negligibly minor impact in the clustermap too.

This ordering has direct practical implications for experimental planning: approximately one third of the variation in harmonization quality is attributable to the method and a further quarter to the batch pre-selection. Importantly, this analysis argues against the assumption that imputation strategy is a critical hyperparameter: we show that both by R^2^ (Figure 6B) and visually by clustermap (Figure 3) imputation is the third factor by importance out of the total four ones.

We then visualized the clustermap best approaches along the six axes of local and global performance by batches and biology (Figure 6C, D). The approaches performed best in global batch overlapping (0.65 – 1.0), followed by local biology preservation (0.07-0.7). Global biology preservation and local batch mixing were the worst performing axes (Figure 6C). The SVA best approaches behaved better in terms of local batch mixing, whereas FSQN R had better biology performance. Among batch removal strategies, C showed the highest batch mixing by kBET and the best biology separation (Figure 6D): the top C approach had 0.43 average batch mixing by kBET and 0.49 average global biology separation. Local biology separation was the best in the J strategy (0.69), and the S0 best approaches were the worst ones by all metrics except for global batch mixing (1.0 in the best S0 approach) and local biology separation (0.55 in the best S0 approach).

We then assessed the contribution of each metric type to the composite score, again for the 2,234 successful approaches. The stacked metric-type composition (Figure 6E) revealed that the top-ranking methods derived their advantage primarily from local neighborhood metrics and distributional similarity metrics: the local ones contributed 0.11 to the clustermap best group and 0.034 to the worst, 27_dwd (0.035 in 14_qsmooth, the second poorest one and the worst by global and local metrics), and the distributional ones added 0.076 to the clustermap best ones, 0.049 to 27_dwd and 0.053 to 14_qsmooth. In contrast, most methods showed comparable performance on global distance metrics: the top 15 methods + clustermap best approaches had them 0.328 (23_vst) to 0.360 (the clustermap best), 0.276 for 27_dwd and 0.275 for 14_qsmooth.

The two top methods after the clustermap best group (it was the best by the composite score with 0.602) were 22_tmm and 23_vst (0.589 and 0.583); both were applied only to strategy C, which limits their comparability with methods run across all 14 strategies (Supplementary Figure 6). The third method was 33_amdbnorm, which completed only 14 of its 84 possible runs, again limiting its comparability with the rest. 28_npn and 10_mnn were the 4^th^ – 5^th^, respectively, with gradual decrease of local composite impact from 0.083 to 0.072 (in 28_npn) and then rising to 0.106 in 10_mnn, accompanied by a decrease in global and distributional performance. The high relative contribution of local and distributional metrics explains the poor discriminative power of rankings based on PCReg or the dispersion separability criterion (DSC) alone — which dominated early-generation harmonization benchmarks (Leek et al. 2010) — because the most biologically consequential improvements are captured only by the expanded 87-metric framework.

It should be noted that the composite score is comparable only between the methods, not between strategies. Because annotation columns become invariant in single-platform or single-biomaterial subsets, the number of defined metrics per approach varies between the strategies (Supplementary Figure 6) and only 1,354 of the 2,234 approaches (60.6%) have all the 87 scoring metrics.

### 3.5. Best approaches assessment by biomarker expression preservation and biology prediction quality

In order to independently assess harmonization quality, we compared metrics of biomarker genes correlation and rank preservation (Supplementary File 2, sheet Table_S3_gene_panel), as well as FL/DLBCL prediction quality for the best approaches and their entire set (Supplementary Figure 18) and visualized the within-biology groups PCA and tSNE plots for ABC and GCB DLBCL subtypes before and after harmonization (Supplementary Figure 19).

Biomarker gene correlation before and after harmonization, averaged by cohorts, was near 1 for the linear methods but as low as -0.057 to 0.058 for the MNN clustermap best approaches (Supplementary Figure 18A), indicating that the high visual performance is achieved at the cost of biology corruption. The rest methods showed the correlation above 0.7 which was considered as acceptable level of biomarker genes preservation. The control correlation over housekeeping genes showed no difference between the clustermap best approaches and rest approaches, calculated using the extended set of 15 genes or the PGK1 gene that was present in all combinations of batch removal strategies and imputations (Supplementary Figure 18B, C). Cross-batch agreement of biomarker genes in the same biology groups was 0.789 median for the clustermap best approaches and 0.627 for the rest approaches (Mann-Whitney two-sided p = 2.6 * 10^-5^, Supplementary Figure 18D), whereas correction by the same agreement for different biology and for the initial non-harmonized expression matrices (Supplementary Figure 18E, F, respectively) revealed no significant difference between the best and the remaining approaches.

To measure cross-batch prediction quality, we calculated leave-one-batch-out (LOBO) macro F1 score and AUC averaged over the batches for the FL versus DLBCL and the three classes (FL, DLBCL and normal B cells) prediction and found no significant difference between the clustermap best approaches and the remaining approaches (Supplementary Figure 18G for F1 in three classes and Supplementary Figure 18H for AUC in two classes). LOBO F1 score in the clustermap best approaches had median value of 0.629 and the highest value 0.877 for SVA in RNA-seq only, Softimpute, whereas the remaining approaches had median value of 0.717. Due to the imbalanced classes, AUC in 2 classes had median 0.913 in clustermap best approaches and 0.903 in the remaining ones.

We then compared biomarker rank concordance in same versus different biology groups without and with a control against the raw datasets (Supplementary Figure 18I, J, respectively) and we found strong linear correlation in both cases (Spearman correlation 0.976, 0.967, respectively, p < 10^-200^), with clustermap best approaches occupying right upper part of the plots, indicating that if a harmonization method converges biomarker gene profiles for the same biology group, it converges them for different groups to the same extent. As an exception, three 08_inmoose_combatseq and 06_combat_seq FFPE only approaches were located above the main diagonal, suggesting that they correctly preserve the biology with lower than usual biology corruption in different groups. It should be noted that MNN harmonization, completely losing the biomarker correlation before and after it, converged gene sets from same and different biology to the same ranking (Spearman correlation 1, Supplementary Figure 18D, 18K), and the same was true for 12_scanorama approaches – rendering these methods useless for the biology-preserving harmonization.

We separately investigated prediction quality by RNA batches (Supplementary Figure 18L) and controlled it over a set of biological group labels being permuted 100 times (Supplementary Figure 18M). We derived that prediction quality was highly heterogeneous by batches: F1 score of the 3 classes LOBO of small microarray batches (GPL20188_FF, GPL17077_FF, GPL16686_FF, GPL23541_FF, GPL13158_FF) have peaks and medians at 1 for all approaches but are shifted towards the 0.6-0.8 levels (batch wise medians 0.625 – 0.822) for the clustermap based approaches, albeit only for the batch GPL16686_FF differences were significant (raw Mann-Whitney two-sided p = 0.021). Simultaneously, large RNA-seq, GPL570 and GPL14951 batches showed comparable performance (Mann-Whitney two-sided p > 0.05). For the permutation control, as expected, randomly permuted LOBO values were in the low value interval of median 0.273 (median absolute deviation 0.014) for the clustermap best approaches, and median 0.241 (median absolute deviation 0.033, Mann-Whitney two-sided raw p = 0.0046).

As a final quality check, we visually investigated DLBCL activated B cell (ABC) and germinal center B cell (GCB) cell of origin (COO) subtypes separation before and after the clustermap best harmonization (Supplementary Figure 19). SVA in RNA-seq did not improve separation between the COO subtypes (Supplementary Figure 19A), whereas MNN, surprisingly, preserved asymmetrical distribution of ABC/GCB in the larger batch groups compared to the isolated batches before harmonization, as can be seen in tSNE and not in PCA plot (Supplementary Figure 19B). FSQN R preserved this co-association worse in the full S0 dataset (Supplementary Figure 19C). The same trend of good ABC/GCB separation for MNN and moderate one for FSQN R was observed for the FF only strategy (Supplementary Figure 19D), whereas SVA in FFPE-only did not form a single connected ABC or GCB groups within the DLBCL multiplatform large group (Supplementary Figure 19E). Finally, AMDBNorm and FSMVN both showed moderate level of COO subtypes preservation (Supplementary Figure 19F, G, respectively).

Taken together, the additional assessment of the clustermap best approaches by biomarkers preservation and prediction quality demonstrated that the MNN approaches achieve high performance at the cost of complete biology corruption, and the remaining best ones do preserve biology (especially FSQN R) but do not confer significant improvement in prediction quality.

### 3.6. Final decision tree for bulk transcriptomic datasets harmonization

Based on the comprehensive benchmarking analysis, we formulated a data-driven hierarchical decision tree for harmonization method selection, conditioning recommendations on dataset biomaterial type and platform composition (Figure 6F). The decision tree formalizes five primary scenarios derived from the benchmark results: 1) fresh-frozen only (FF) with RNA-seq and microarrays, 2) FFPE only with RNA-seq and microarrays, 3) RNA-seq only (any biomaterial including FF and FFPE), 4) RNA-seq combined with Illumina microarrays (any biomaterial), and 5) the full dataset: RNA-seq combined with microarrays with any biomaterial. We did not include the trivial combination of single platform and biomaterial type (such as FFPE-only RNA-seq, or FF-only Affymetrix), because such datasets already co-clustered in PCA and tSNE space (Figure 4A), so extensive harmonization was not required. We considered only the cross-platform and/or cross-biomaterial datasets as the most challenging ones.

For fresh-frozen only datasets — comprising Affymetrix, Illumina, and Agilent microarray platforms mixed with RNA-seq — MNN achieved the best overall performance across all metric families but was excluded because the biomarker correlation vanished, so FSQN R is recommended. For FFPE-only datasets, SVA with Softimpute or KNN imputation represents the rarely demonstrated successful mixing of FFPE Affymetrix, FFPE Illumina microarray, and FFPE RNA-seq platforms in a single harmonized GC B-cell lymphoma dataset; this combination achieved a 56% relative reduction in PC1 variance (from 87.6% raw to 38.4% post-harmonization, Supplementary Figure 15) while preserving FL and DLBCL cluster separation. For RNA-seq only datasets, SVA with Softimpute or KNN again emerged as the top-performing approach.

For mixed RNA-seq and Illumina microarray datasets, FSQN R achieved the best balance between global correction and biology preservation, with AMDBNorm or FSMVN combined with post-removal as alternatives for scenarios where global PCReg is prioritized. For the most compositionally complex scenario — RNA-seq combined with Affymetrix and Illumina microarrays spanning multiple biomaterial types — FSQN R is proposed instead of MNN, despite the latter achieving superior global and local batch correction at the cost of biomarker rank correlation being lost. A common finding across the RNA-seq only and FFPE only scenarios was the superiority of imputation-augmented approaches over strict gene-set restriction when combined with SVA harmonization (Figures 4 and 5).

Notably, post-removal — the exclusion of one PCA-outlier batch after harmonization — provided marginal metric improvement (R² < 0.01 in the variance decomposition), confirming that it is a fine-tuning step rather than a fundamentally distinct correction stage. We recommend post-removal as an optional step for all strategies, applied after visual inspection (Figure 6F) when a strong outlier batch is consistently identified across multiple imputation conditions.

## 4. DISCUSSION

### 4.1. Subtle biological differences under strong batch effect

We aimed to find the best harmonization approach for the multiplatform GC B cell lymphoma and normal B cells bulk transcriptomic dataset of 7,174 samples. We assembled it from bulk profiles since cancer bulk transcriptomics still outnumbers single-cell-profiled patients by more than an order of magnitude (Zhang et al. 2021; Cho et al. 2026) (Liu et al. 2025; Zeng et al. 2025) and is routinely used in clinical assays such as BostonGene Tumor Portrait (Yudina et al. 2025) and FoundationOneRNA (D. Sun et al. 2025). Yet the analogous approach can be applied for scRNA batch correction, albeit we anticipate different challenges there, as the dominant FF/FFPE batch effect is largely irrelevant to scRNA-seq. In the future FL studies, our research consortium plans to include single cell datasets too.

The dataset assembled for this study was markedly imbalanced with respect to both batch and biology group composition (Figure 1D, 1E), reflecting the typical structure of publicly available retrospective transcriptomic data aggregated from independent clinical studies. This real-world heterogeneity — with dominant DLBCL-only batches, rare subtypes confined to one or two cohorts, and mixed FF/FFPE biomaterial composition — renders the present FL-containing dataset an inherently more challenging harmonization target than purpose-built benchmark datasets (Borisov and Buzdin 2022).

Moreover, the follicular lymphoma transcriptome represents one of the most technically demanding targets for cross-platform harmonization precisely because the biological signal of interest — the distinction between FL and DLBCL, or between FL molecular subtypes — is subtle relative to the technical noise introduced by inter-platform and inter-biomaterial differences (Figure 4A). Our dataset, spanning 7,174 samples across 29 RNA batches and four transcriptomic platforms, encapsulates this challenge at a scale not previously assembled for FL research. The observation that 77.4% of PCA variance in the raw S0 dataset is attributable to RNA_BATCH — without biology-driven structure visible before harmonization — quantifies the magnitude of this challenge and underscores why standard benchmarking frameworks using TCGA or GTEx-scale datasets, where biology effects typically dominate batch effects, are insufficient for subtle-biology applications (Hicks et al. 2018; Borisov et al. 2022; Skubleny et al. 2024).

Current benchmarking datasets (TCGA, GTEx, SEQC, MAQC etc) comprise biologically highly distinct groups, such as pan-cancer models comparing brain, colon, and lung tissues, coupled with relatively minor technical variations, such as homogeneous RNA-seq protocols derived from high-quality fresh-frozen tissue (Lonsdale et al. 2013; Weinstein et al. 2013). If an algorithm utilizes a standard location-and-scale shift, such as simple mean-centering, or forces all sample distributions to match a theoretical reference via standard quantile normalization, it will successfully eliminate these minor batch effects while preserving thousands of differentially expressed genes between colon and brain (Hicks et al. 2018). However, the vast majority of clinically actionable biomarker research (Sorokin et al. 2020; Zottel et al. 2020; Vladimirova et al. 2021b) does not attempt to distinguish a brain tumor from a colon tumor; rather, it attempts to distinguish closely related disease states, therapeutic prognoses, or subtle developmental stages within a single lineage (Sorokin et al. 2021; Gudkov et al. 2022b; Sorokin et al. 2022; Kang et al. 2023). Our analysis shows that among the 31 harmonization tools analyzed, only SVA and FSQN R preserved the subtle differences between FL, DLBCL and normal GC B cells under strong batch effect, both globally and locally, allowing for transcriptomic biomarkers mining. FSMVN and AMDBNorm showed promising performance under a specific context of RNA-seq and Illumina microarrays combined.

The FL-specific biological discovery — two transcriptional subgroups revealed by SVA with Softimpute in the RNA-seq-only strategy — constitutes an unexpected and biologically significant finding emerging from the harmonization process itself. These subgroups were invisible in all other harmonization configurations, including other methods in the same strategy and SVA with strict imputation. This dependency on imputation suggests that the subgroup signal resides in genes recovered by Softimpute beyond the strict 3,447-gene set, genes that may carry platform-specific regulatory signatures detectable only at higher coverage depth. Whether these subgroups correspond to the established GCB-like and memory-cell-like (Laurent et al. 2024) molecular subtypes of FL or represent a novel transcriptional axis orthogonal to known classifiers, requires downstream survival analysis and gene set enrichment that is beyond the scope of this benchmarking study. Crucially, the finding underscores that harmonization method selection is not merely a technical pre-processing decision but can determine which biological discoveries are accessible from a given dataset.

### 4.2. Novel computational pipeline for harmonization tools adjustment and selection

The ComboBatch pipeline is conceptually distinct from previous harmonization benchmarks (Leek et al. 2010; Borisov and Buzdin 2022; Yu et al. 2024) in three fundamental respects. First, it exhaustively evaluates the full cross-product of four hyperparameter dimensions — 14 batch removal strategies × 3 imputation methods × 33 harmonization methods × 2 post-removal conditions = 2,772 theoretical configurations (2,407 completed, 2,234 successfully completed) — instead of selecting a representative subset of configurations for pairwise comparison. This exhaustive coverage allows the pipeline to detect interactions between factors (such as the imputation-dependency of SVA’s FL subgroup discovery) that would be missed by incomplete designs. Second, it applies 87 polarity-defined scoring metrics spanning local neighborhood, global distance, distributional similarity and other aspects of harmonization quality, providing a multi-dimensional evaluation surface rather than relying on a single metric or a small panel. Additionally, prediction quality and biomarker correlation metrics are applied as an independent evaluation, showing that MNN harmonization attained the best performance with the cost of biologically relevant correlations lost. Third, it operates under the constraint of realistic biological subtlety: the FL/DLBCL distinction is ten-fold smaller in transcriptional effect size than the batch effects in this dataset: PCReg by batch is 0.226 (fraction of variance explained by batch 0.774) and by biology is 0.923 (fraction of variance explained 0.077) in the S0, 01_raw, strict and no post-removal dataset (Supplementary File 3), requiring approaches that simultaneously correct for large technical noise while preserving small biology signals.

The computational scale of ComboBatch — 2,234 successfully completed approaches on 32–48 vCPU / 240 GiB RAM infrastructure with parallelized job dispatch across S3-backed intermediate storage — required systematic engineering of timeout handling, memory-limited execution, and error recovery. All harmonization methods that exceeded 3 hours of CPU time were recorded as failures and excluded from the main analysis (see Methods and the repository https://github.com/Nikit357/FL_harmonization/tree/main/harmonization-scripts). The pipeline architecture is containerized and reproducible, with intermediate expression matrices for public cohorts and metric tables available via the Zenodo links described in the Data Availability statement.

We note important technical limitations of the current pipeline. The kBET metric is known to be sensitive to small batch sizes and failed to produce valid outputs for strategies with very small batch representations (Büttner et al. 2018), and the present dataset contains 2 batches and 15 cohorts with fewer than 10 samples. Two of the six methods that failed implementation, 30_recombat and 37_fabatch, did so for technical reasons that remain to be resolved. Moreover, the field of transcriptomic harmonization tools is expanding rapidly, including recent diffusion-model-based methods (Cui et al. 2025), so the current review of 75 methods should be revised periodically, which we plan to do in the upcoming GitHub ComboBatch releases (the repository is available via the link https://github.com/Nikit357/ComboBatch).

We also acknowledge additional mathematical limitations of the ComboBatch pipeline that require further improvements in the next releases. First, although the composite score was not a decision criterion, it averages different metric subsets for different strategies – rendering them incomparable. Second, six of the 31 analyzed methods were not run on all 14 strategies, adding another layer of complexity into the comparison logic. Third, the embedding-space metrics were computed on stochastic embeddings without replication – averaging over multiple repeats could be an improvement here.

Finally, the overall study design could be improved by applying the LOBO principle to selection of the best approaches: identifying the best approach for a dataset with 2-5 RNA batch groups removed and then testing the approaches selected on the batches excluded. Such design could test robustness and transferability of the current approach, albeit prediction quality metrics with random permutation already partially assess it.

### 4.3. Clustermap on a full metrics space as the emerging approach for harmonization selection

The 87-metric × 2,234-approach clustermap (Figure 3) with additional prediction quality and biological correlation assessment (344 total metrics) represents, to our knowledge, the largest systematic harmonization evaluation matrix reported to date in bulk transcriptomics. The clustermap revealed a robust four-cluster structure at the approach level — good cross-platform, bad cross-platform, same sample type, and same platform — that was reproduced across independent Spearman correlation analysis (Supplementary Figure 9) and metric-space embeddings (Supplementary Figure 10). Critically, this structure was dominated by batch removal strategy instead of harmonization method (as the R^2^ analysis in Figure 6B), confirming the theoretical expectation that sample composition determines the accessible range of correction.

The practical value of the clustermap-based selection approach is that it provides a data-driven ranking that does not require prior assumptions about which metrics are most relevant for a given dataset. For datasets with different biological contexts — rare cell types, subtle molecular subtypes, or extreme platform imbalance — the same pipeline can be run with the same 87 metrics, and the clustermap will identify the dominant sources of quality variation specific to that context. The ability to visually identify anomalous metric-level patterns — such as the HarmonizR gene-retention failure (manifesting as a distinct band in the NA-percentage metric cluster in Figure 3) — provides a level of interpretability that single-score rankings cannot achieve.

The metric cross-correlation structure (Supplementary Figure 7) further reveals that the local and global metric clusters behave in an orthogonal manner (Spearman r median 0.009 and MAD 0.20 between clusters), supporting the two-axis evaluation framework used throughout the benchmark. This empirical orthogonality is the key statistical justification for requiring both metric families in harmonization benchmarks, and it explains why methods that score highly on PCReg can simultaneously perform poorly on kBET and iLISI.

It should be noted that even 87 scoring metrics clustermap was shown susceptible to errors: MNN, being the best approach by the clustermap metrics, was demonstrated to corrupt rank structure of biomarker genes (Supplementary Figure 18) and was excluded from the best approaches. Paradoxically, it preserved the COO structure in tSNE plots (Supplementary Figure 19) better than the final clustermap best approaches. Taken together, these examples highlight the necessity of integrated evaluation workflows of transcriptomic harmonization, that should include cross-validation, quality metrics calculation and expert visual inspection.

### 4.4. AI-assisted methods evaluation made the same impact as the expert-driven method discovery

The ComboBatch benchmark was designed as an exhaustive computational search which revealed that of the two top-performing methods, SVA and FSQN R, the former was identified through AI-assisted literature search and method curation (Liu et al. 2026 Apr 14), while the latter was initially discovered through manual literature exploration of the quantile normalization family.

Interestingly, rank normalization and standard quantile normalization — both harsh and computationally simple approaches — performed substantially below their more architecturally advanced FSQN R counterpart. This performance gap cannot be explained by harshness alone: FSQN R shares the quantile normalization principle but matches each feature to a gene-wise reference distribution derived from a reference platform, preserving expression shape while eliminating scale differences (Skubleny et al. 2024). In this dataset, methods that condition on biological structure or model latent factors outperformed generic location-and-scale corrections, even when the simpler methods appear algorithmically equivalent at first inspection (Yu et al. 2024).

### 4.5. Comparison with other batch effect mitigation approaches

Among the 31 methods analyzed, SVA and FSQN R were the only algorithms that simultaneously resolved FL, DLBCL, and normal GC B-cell transcriptional differences after cross-platform integration, with AMDBNorm and FSMVN having a narrower scope of application. This finding is consistent with the known mechanistic properties of these methods: SVA uses surrogate variable analysis to model latent technical factors while conditioning on the provided biological covariates (Leek and Storey 2007) and FSQN R applies feature-specific quantile normalization against an RNA-seq reference distribution, preserving the global expression topology of the reference biology (Franks et al. 2018). Both methods share a common architectural principle: they model or condition on biological structure during the correction process, preventing the collapse of biology-associated variance that characterizes unconstrained normalization methods.

Methods that dominated earlier harmonization benchmarks designed on TCGA or GTEx data — pyCombat (Behdenna et al. 2023), limma removeBatchEffect (Ritchie et al. 2015), median scaling (Kappal and https://independent.academia.edu/SunilKappal 2019 Jan 1), and standard quantile normalization (Qiu et al. 2013) — achieved strong global batch correction (PCReg reduction) in our dataset but failed both to preserve local biology structure and to mix batches locally (Figure 3). This failure is directly attributable to the scale mismatch between training context and our dataset: in TCGA or GTEx, batch effects are typically 2–5-fold smaller in variance explained than in our multi-platform lymphoma dataset (Lonsdale et al. 2013; Weinstein et al. 2013), allowing unconstrained methods to correct batch effects without inadvertently erasing biology. When the technical noise is ten-fold larger than the biology signal — as in our raw FL dataset — unconstrained methods cannot distinguish the two and remove both.

The Quartet reference material framework (Yu et al. 2023), while conceptually elegant for prospective experimental designs, is inapplicable to retrospective harmonization of pre-existing clinical transcriptomic cohorts such as ours, where concurrent profiling of a reference material was not performed. Our results suggest that for retrospective multi-platform integration under realistic clinical-cohort conditions — heterogeneous batch sizes, multiple platforms, mixed biomaterials, confounded batch–biology composition — algorithm selection conditioned on biomaterial type and platform composition, as formalized in the ComboBatch decision tree, provides a robust and reproducible alternative to reference-material-based normalization frameworks. The decision tree further provides actionable guidance for prospective study design by identifying which sample composition scenarios are most tractable and which harmonization approaches are most robust to deviations from those scenarios.

## 5. CONCLUSIONS

In the present study we assembled a GC B cell lymphoma cross-platform cohort of 7,174 bulk transcriptomic profiles. We built the ComboBatch pipeline and showed that among the 31 harmonization tools analyzed, only SVA and FSQN R reproducibly resolved subtle differences between the transcriptional programs in FL, DLBCL and normal GC B cells under strong batch effects between RNA-seq and microarrays, as well as between FFPE and FF biomaterials. FSQN R was the second priority method, and AMDBNorm and FSMVN could be applied for harmonization of RNA-seq and Illumina microarrays. The 33 harmonization and 3 imputation methods implemented in the study are available as a versatile ComboBatch tool with the Docker image that contains all the R and Python libraries installed. We hope that the tool and the real-world benchmarking scenario under the subtle biology and strong batch effect will be applicable both in the B cell lymphoma and cancer transcriptomic biomarkers mining fields.

## 6. ACKNOWLEDGEMENTS

We thank Alisa Sadekova for normal B cells dataset provision and Alexander Ryabykh for assistance with open-source datasets gathering. We deeply appreciate support and fruitful discussions with Katerina Nuzhdina, Aleksei Efremov, Konstantin Chernyshov, Daniil Ivanov and Anastasiya Yudina. D.N. thanks his family — his sons Ivan and Alexandr and his wife Irina Nikitina — for their patience and support during the preparation of this manuscript.

## 7. AUTHOR CONTRIBUTIONS

Daniil Nikitin: Conceptualization, Data curation, Formal analysis, Investigation, Methodology, Software, Validation, Visualization, Writing — original draft, Writing — review & editing. Nikolay Borisov: Conceptualization, Methodology, Writing — review & editing. Maria Savchenko: Conceptualization, Methodology. Anatoly Bobe: Data curation, Writing — review & editing. Mark Meerson: Conceptualization, Resources, Data curation, Software. Alexander Nesmelov: Data curation. Nazar Harutyunyan: Data curation. Svetlana Paponova: Data curation. Andrey Kravets: Data curation. Alexandr Zaitsev: Writing — review & editing. Alexandr Bagaev: Conceptualization, Supervision, Resources, Funding acquisition. Arsen Arakelyan: Conceptualization, Supervision, Methodology, Funding Acquisition, Software, Project administration, Writing — review & editing.

## 8. SUPPLEMENTARY DATA

Supplementary Data are available at NAR online.

**Supplementary Figure 1.** (A) Bar plot of sample counts by cohorts and biology, ordered by cohorts. Biology groups are color coded according to the color legend inserted. (B) Bar plot of sample counts by cohorts and batches, ordered by cohorts. Batches are color coded according to the color legend in the panel. (C) PCA plot of an initial FSQN R normalization of the dataset, colored by batches as in the panel B. The arrow shows batches that were excluded for the strategy A.

**Supplementary Figure 2.** Gene expression distribution plots for the S0 strategy with strict NA handling, log2-transformed. Each plot contains gene expression distribution of all samples in a given cohort.

**Supplementary Figure 3.** Gene expression distribution plots for the S0 strategy with KNN imputation, log2-transformed. Each plot contains gene expression distribution of all samples in a given cohort.

**Supplementary Figure 4.** Gene expression distribution plots for the S0 strategy with Softimpute imputation, log2-transformed. Each plot contains gene expression distribution of all samples in a given cohort.

Supplementary Figure 5. Clustermap showing Jaccard index of gene sets intersections between all the batch removal strategy and imputation combinations.

Supplementary Figure 6. Heatmap indicating number of calculated metrics for all combinations of batch removal strategies and imputations (rows) and harmonization levels and post-removals (columns).

Supplementary Figure 7. Clustermap of the 87 scoring metrics Spearman cross-correlations in the space of the 2,234 successful harmonization approaches. Metrics are color annotated by computational group, annotation column and type.

Supplementary Figure 8. 3 X 3 square grid of scatterplots showing distribution of the 87 scoring metrics in PCA, UMAP and tSNE embeddings (columns) calculated on the space of the 2,234 harmonization approaches, and color annotated by metric type (top row), annotation column for metric (middle row) and metric computational group (bottom row).

Supplementary Figure 9. Clustermap showing Spearman cross-correlations of the 2,234 successful harmonization approaches in the space of the 87 scoring metrics. Harmonization approaches are color annotated by their batch removal strategy, imputation, harmonization method, post-removal and harshness level. Major clusters are differentially color coded and named on the top of the clustermap.

**Supplementary Figure 10**. 3 X 5 rectangular grid of scatterplots showing distribution of the 2,234 successful approaches in PCA, UMAP and tSNE embeddings (columns) calculated based on the 87 scoring metrics space. The approaches are color annotated from top to bottom row by batch removal strategy, harmonization method, harshness level, imputation and post-removal.

**Supplementary Figure 11.** (A) 4 X 2 rectangular grid of PCA plots of 10_mnn harmonized S0 strategy with strict imputation, H strategy strict, D strategy strict and 01_raw (no harmonization) S0 strategy strict as a baseline. The top row shows scatterplots colored by batches, the bottom row demonstrates the ones colored by biology according to color palettes from Figure 1. (B) The same 4 X 2 rectangular grid of UMAP plots. (C) The same 4 X 2 rectangular grid of tSNE plots.

**Supplementary Figure 12. (**A) 4 X 2 rectangular grid of PCA plots of 04_sva harmonized C strategy with KNN, Softimpute and strict imputation, as well as 01_raw (no harmonization) C strategy strict as baseline. The top row shows scatterplots colored by batches, the bottom row demonstrates the ones colored by biology according to color palettes in the bottom of the figure. (B) The same 4 X 2 rectangular grid of UMAP plots. (C) The same 4 X 2 rectangular grid of tSNE plots.

Supplementary Figure 13. (A) 4 X 2 rectangular grid of PCA plots of 16_fsqn_r harmonized C strategy with KNN, Softimpute and strict imputation, as well as 01_raw (no harmonization) C strategy strict as baseline. The top row shows scatterplots colored by batches, the bottom row demonstrates the ones colored by biology according to color palettes in the bottom of the figure. (B) The same 4 X 2 rectangular grid of UMAP plots. (C) The same 4 X 2 rectangular grid of tSNE plots.

Supplementary Figure 14. (A) 3 X 2 rectangular grid of PCA plots of 10_mnn harmonized strict, 16_fsqn_r harmonized strict and 01_raw strict J strategy as baseline. The top row shows scatterplots colored by batches, the bottom row demonstrates the ones colored by biology according to color palettes on the left side of the figure. (B) The same 3 X 2 rectangular grid of UMAP plots. (C) The same 3 X 2 rectangular grid of tSNE plots.

Supplementary Figure 15. (A) 4 X 2 rectangular grid of PCA plots of 04_sva harmonized K strategy with Softimpute, KNN and strict imputation, as well as 01_raw (no harmonization) K strategy strict as baseline. The top row shows scatterplots colored by batches, the bottom row demonstrates the ones colored by biology according to color palettes in Figure 1. (B) The same 4 X 2 rectangular grid of UMAP plots. (C) The same 4 X 2 rectangular grid of tSNE plots.

Supplementary Figure 16. (A) 4 X 2 rectangular grid of PCA plots of S0 strategy harmonized with 13_fsmvn with strict, KNN and Softimpute imputation, as well as 01_raw (no harmonization) S0 strategy KNN as baseline. The top row shows scatterplots colored by batches, the bottom row demonstrates the ones colored by biology according to color palettes in Figure 1. (B) The same 4 X 2 rectangular grid of UMAP plots. (C) The same 4 X 2 rectangular grid of tSNE plots.

Supplementary Figure 17. (A) 4 X 2 rectangular grid of PCA plots of 33_amdbnorm harmonized K strategy strict, S0 strategy strict, J strict and J Softimpute. The top row shows scatterplots colored by batches, the bottom row demonstrates the ones colored by biology according to color palettes in Figure 1. (B) The same 4 X 2 rectangular grid of UMAP plots. (C) The same 4 X 2 rectangular grid of tSNE plots.

Supplementary Figure 18. (A) Boxplot comparing mean per-gene Spearman ρ before vs after harmonization (metric name mk_rho_mean_all_genes_narrow_set) for the clustermap best and the remaining approaches, colored by harmonization methods. P is raw Mann-Whitney two-sided p-value, RBC is rank-biserial correlation for Mann-Whitney test used as an effect size estimation (Peres 2025). (B) The same boxplot for spearman ρ before vs after harmonization, biomarker − housekeeping genes (mk_rho_marker_minus_hk_narrow_set). (C) The same boxplot for spearman ρ before vs after harmonization, biomarker − PGK1 gene only (mk_rho_marker_minus_PGK1_only_narrow_set). (D) The same boxplot for cross-batch agreement of biomarker genes of the same biology (xb_rank_agree). (E) The same boxplot for cross-batch agreement of the same biology groups minus the different biology (xb_rank_agree_ratio). (F) The same boxplot for cross-batch agreement by biology, subtracted by non-harmonized approaches (xb_rank_agree_ratio_delta). (G) The same boxplot for leave-one-batch-out macro-F1: FL, DLBCL, normal B-cells (pv_lobo3_f1_macro_mean). (H) The same boxplot for leave-one-batch-out macro-AUC: FL vs DLBCL (pv_lobo2_auc_macro_mean). (I) Scatter plot of cross-batch agreement of extended biomarker genes in the same versus the different biology groups (xb_rank_agree vs xb_rank_disagree_diffbio), colored by harmonization methods and with clustermap best approaches highlighted as black stars. (J) The same scatterplot for cross-batch agreement corrected by the corresponding raw dataset by batch removal and imputation (xb_rank_agree_delta vs xb_rank_disagree_diffbio_delta). (K) Transition plot of cross-batch agreement in same biology groups (xb_rank_agree) between the raw and harmonized datasets, the clustermap best approaches colored by the harmonization method. (L) Violin plot of LOBO macro-F1: FL, DLBCL, normal B-cells by batch and by the clustermap best and remaining approaches. (M) Scatterplot of LOBO macro-F1 3 classes versus the same metric for 100 random permutations of biology labels (pv_lobo3_f1_macro_mean vs pv_lobo3_f1_macro_perm_mean), the clustermap best approaches colored by harmonization methods.

Supplementary Figure 19. PCA and tSNE plots for raw and harmonized datasets, colored by the biology with ABC/GCB COO classification of DLBCL samples. (A) SVA in RNA-seq only KNN versus RNA-seq only raw strict. (B) MNN in S0 no removal strict versus raw S0 strict. (C) FSQN R in S0 no removal strict. (D) MNN and FSQN R in FF only strict compared to FF only strict dataset. (E) SVA Softimpute harmonization of FFPE only dataset compared to raw strict FFPE only. (F) AMDBNorm in S0 no removal strict. (G) FSMVN in S0 no removal.

**Supplementary File 1**. The GC B-cell lymphomas cohort annotation by sample.

**Supplementary File 2.** Excel workbook with five sheets: ‘Genes_per_sample’, the number of genes with defined gene expression for each of the 7,174 samples in the initial dataset prior to imputation; ‘Metric_polarity’, the polarity coefficients, classification and explanation of the 344 harmonization quality metrics used in this study; ‘Table_S1_methods’ (Supplementary Table S1), the 39 harmonization methods reviewed for implementation, with the parameters used, the harshness tier and its justification, and the full bibliographic reference for each method; and ‘Table_S2_metrics’ (Supplementary Table S2), the definitions, types, computational groups and sources of the harmonization quality metric families; and ‘Table_S3_gene_panel’ (Supplementary Table S3), the 633 biomarker genes across 63 signatures with their coverage, quality-control and per-gene correlation statistics.

**Supplementary File 3**. Values of the 87 scoring metrics for the 2,407 harmonization approaches computed in this study; the 2,234 approaches used in the analysis are those with pct_samples_allNA < 5. The full metrics table can be found in the Zenodo record described in the Data Availability statement.

## 11. DATA AVAILABILITY

The data supporting this article are available in the article and its online supplementary materials. Intermediate analysis files, Python scripts, YAML configuration files for the computational pipeline, Jupyter notebooks, and exploratory plots not included in the article or supplementary materials have been deposited in a public GitHub repository (https://github.com/Nikit357/FL_harmonization) and archived at Zenodo (Nikitin). The ComboBatch transcriptomic harmonization tool is available in a separate GitHub repository accessible via the link (https://github.com/Nikit357/ComboBatch), archived at Zenodo (Nikitin). Containerized and parallelized Shambhala-2 that was developed and tested in this article has been published in a separate GitHub repository and is accessible via the link (https://github.com/Nikit357/Shambhala2_fast), again archived at Zenodo (Nikitin).

All the repositories are available for peer review and archived with a permanent DOI at Zenodo.

The transcriptomic data analyzed in this study derive from publicly available and proprietary germinal-center B-cell lymphoma cohorts; their repository accession numbers (GEO / ArrayExpress / SRA) are listed in Supplementary File 1. Processed and raw expression matrices, as well the harmonization metric tables generated for this study have been deposited in a public repository issuing a permanent DOI and are accessible via the link https://zenodo.org/records/22737294 (Nikitin et al.).

