## Supplementary figures and images for "Benchmarking of bulk transcriptomic harmonization tools in a multi-platform B-cell lymphoma cohort identifies feature-specific quantile normalization and surrogate variable analysis as top-performing methods"

### Supplementary Figure 18

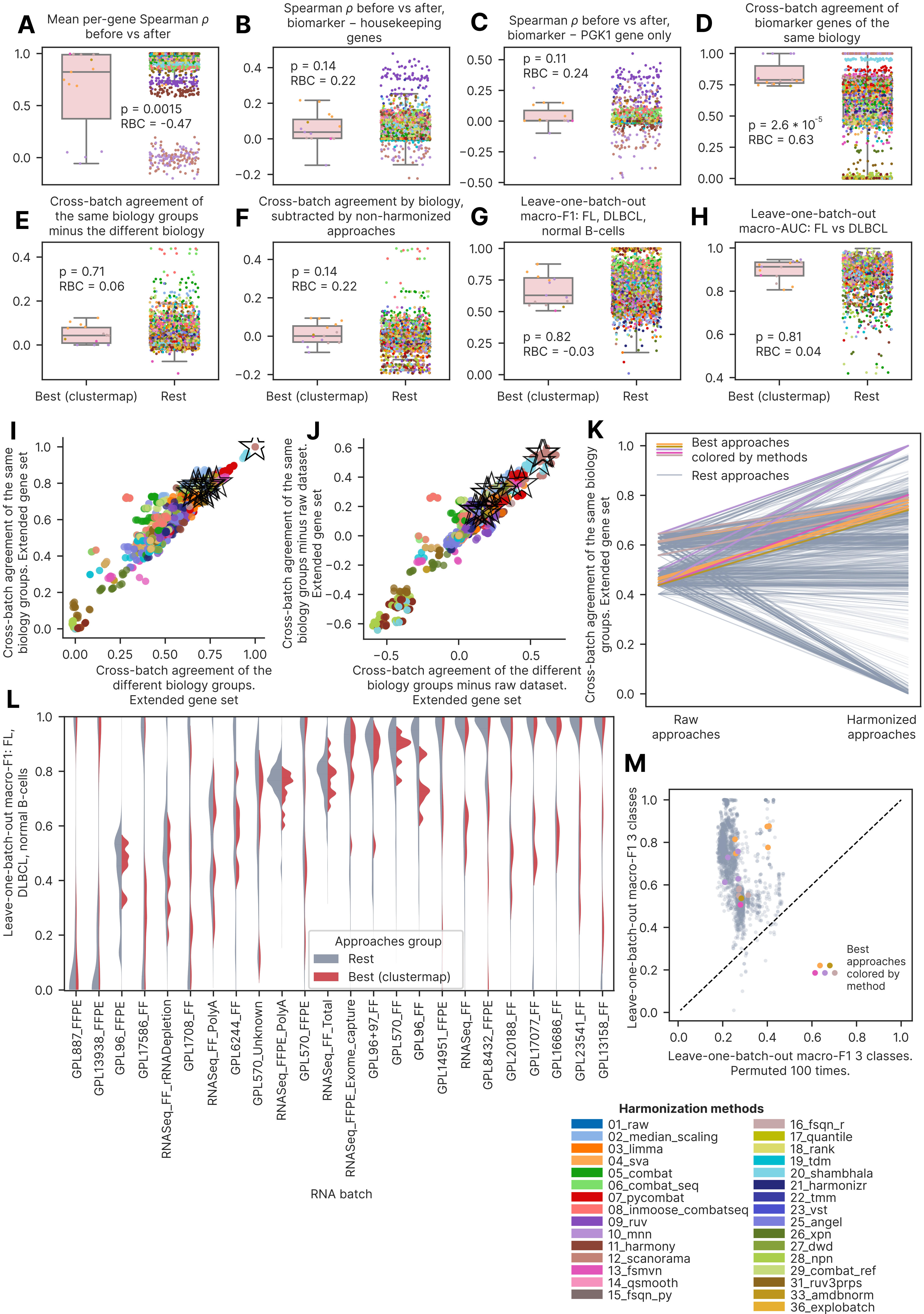

### Supplementary Figure 19

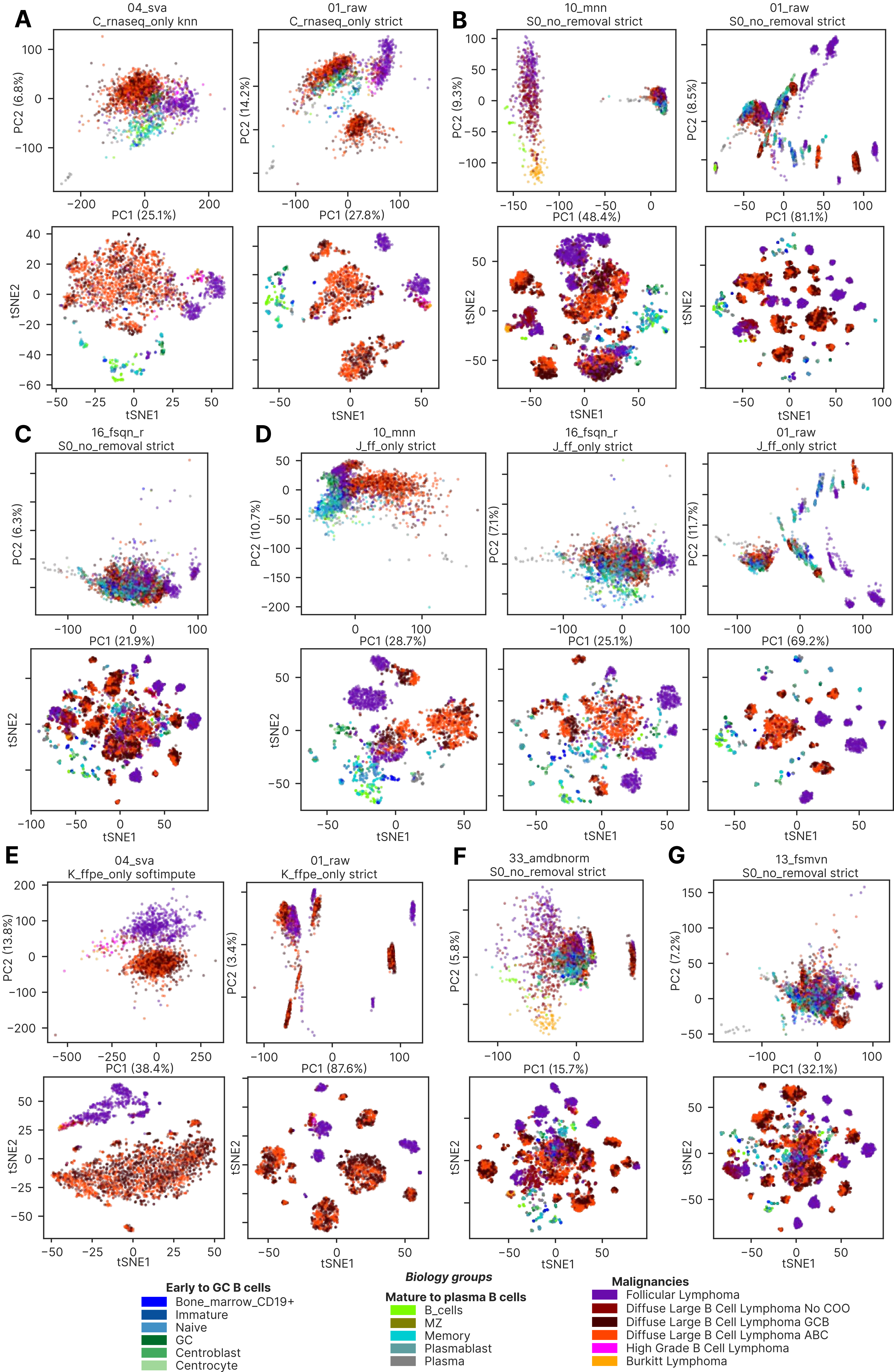

### Supplementary Figures 1-2

Supplementary Figure 1

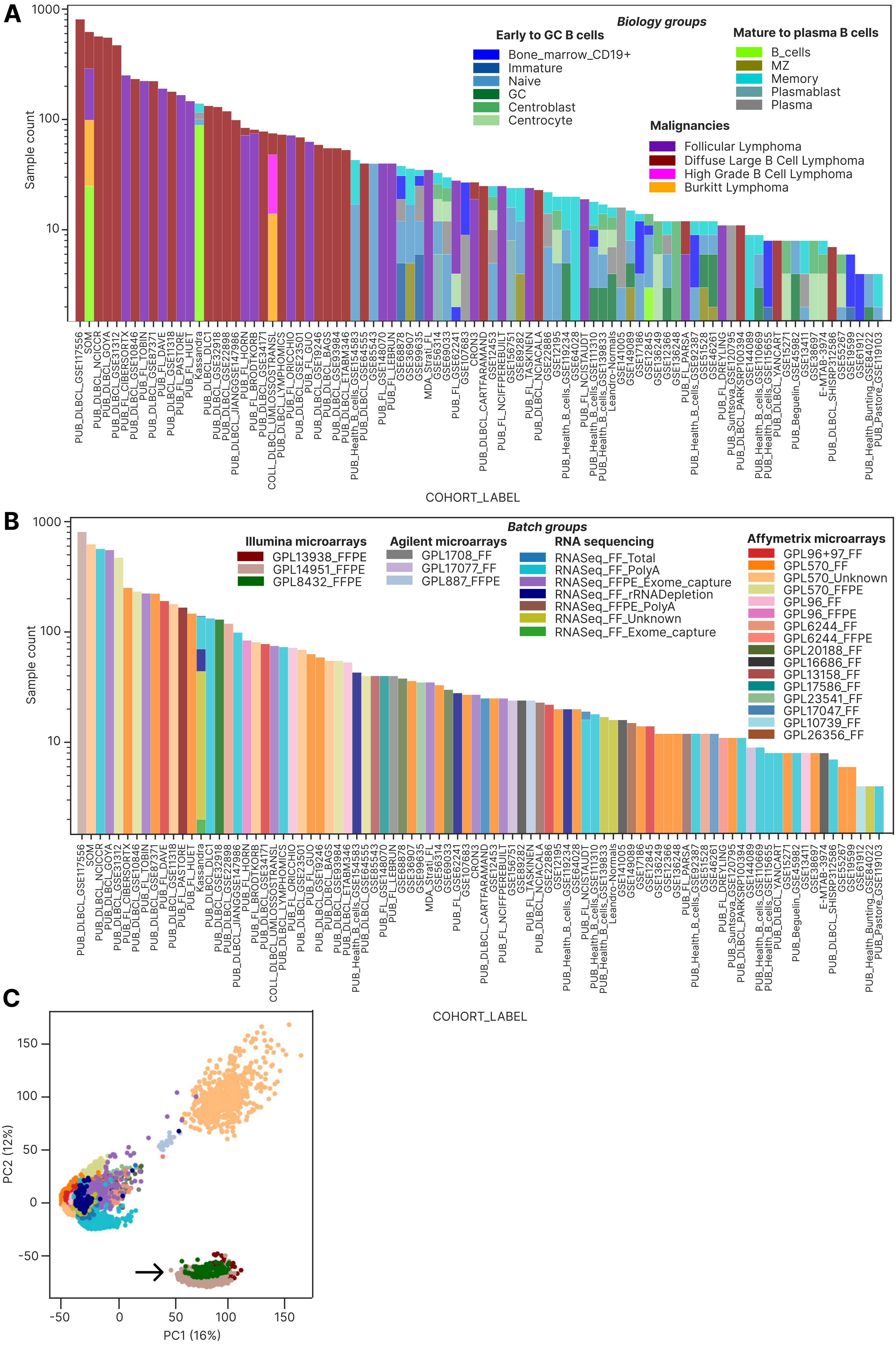

Supplementary Figure 2

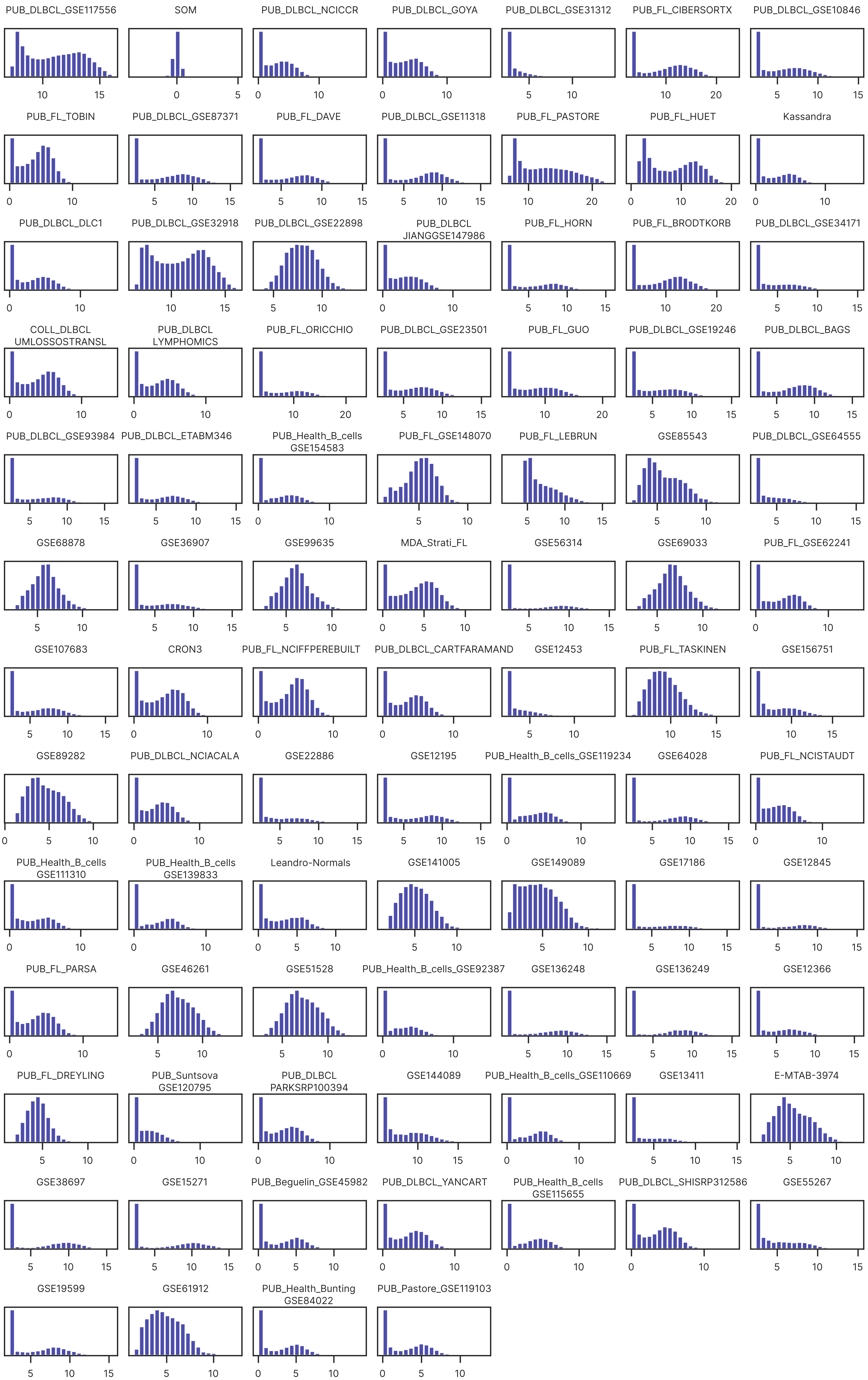

### Supplementary Figures 3-4

Supplementary Figure 3

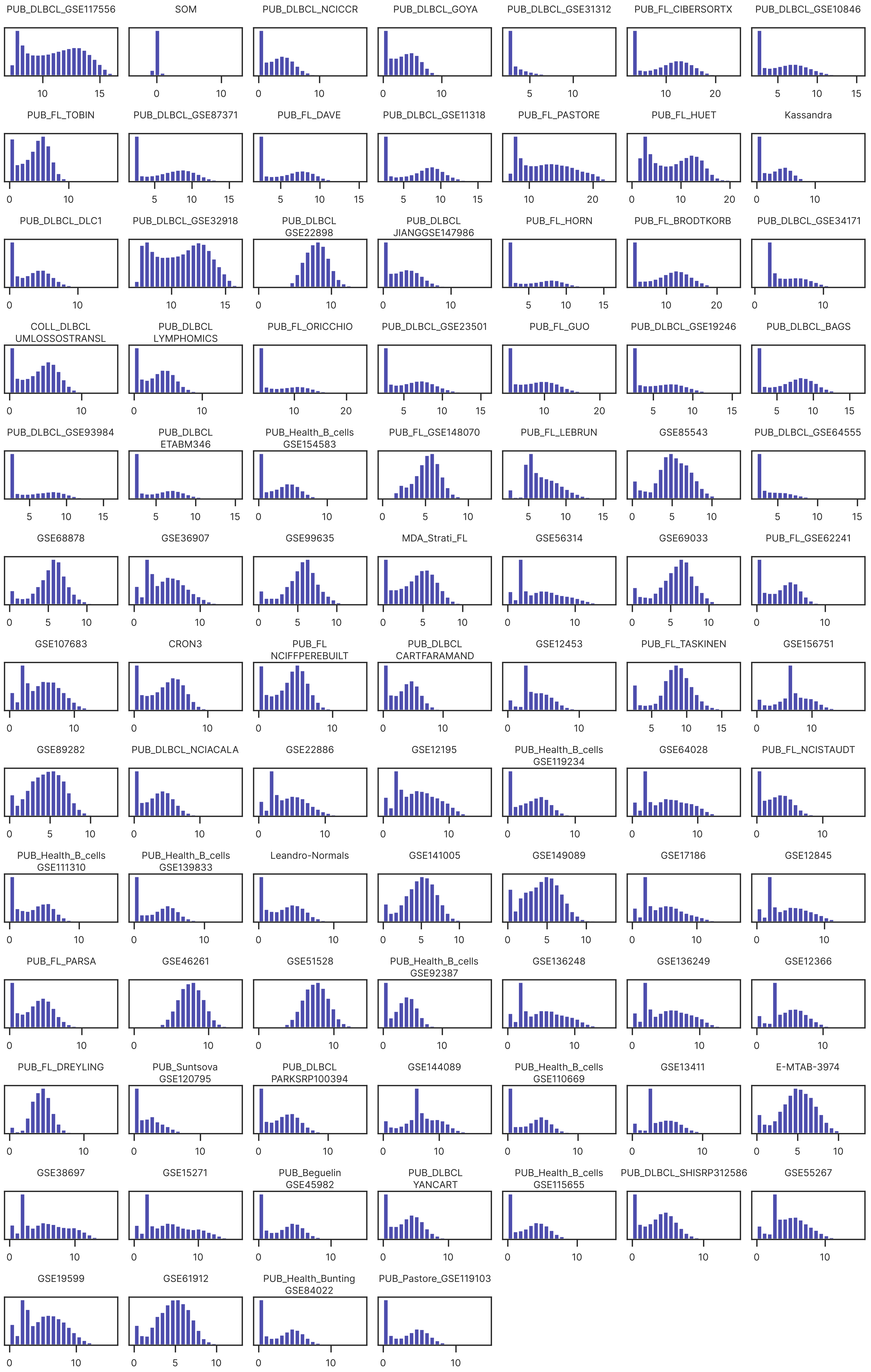

Supplementary Figure 4

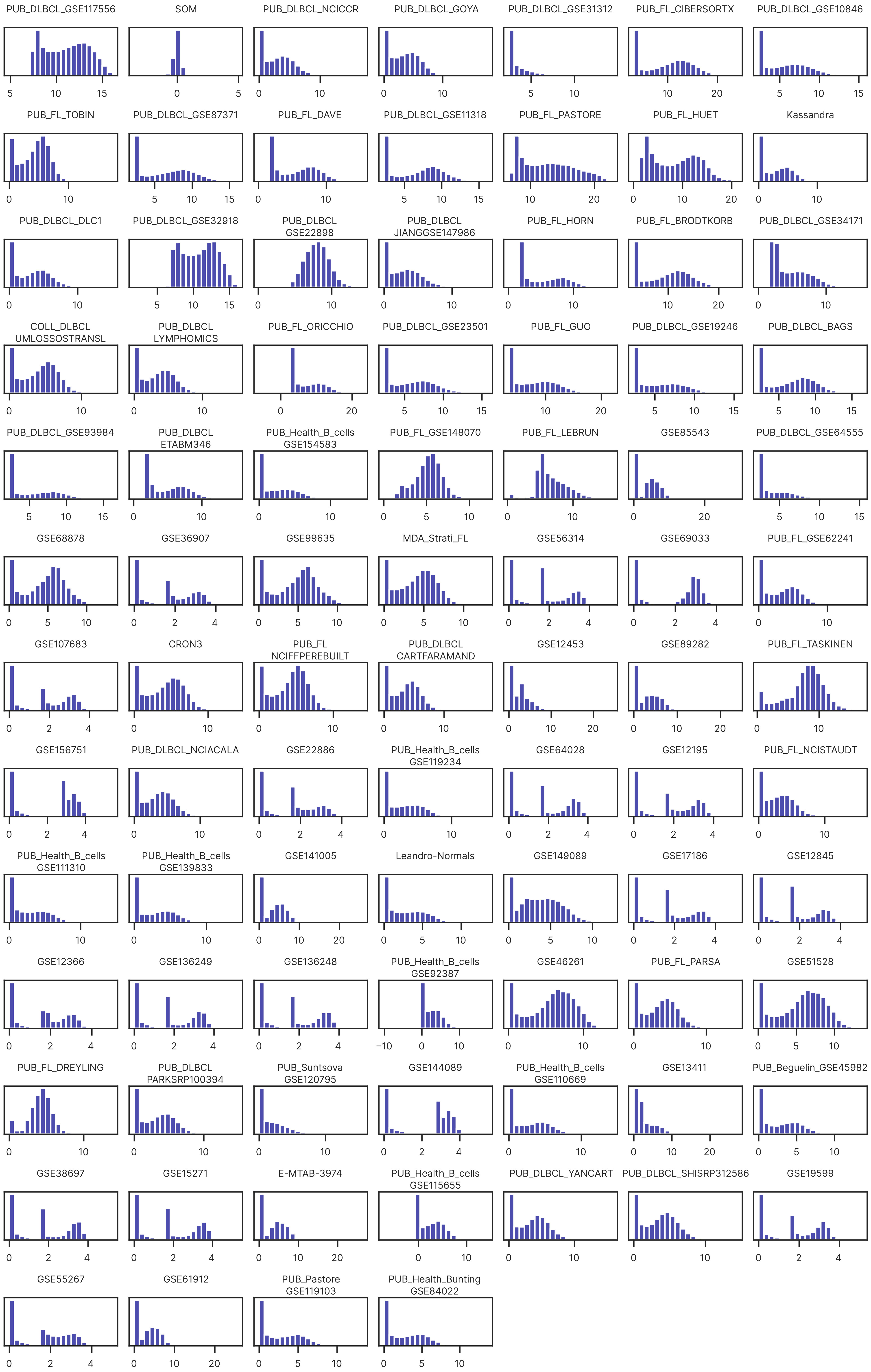

### Supplementary Figures 5-7

Supplementary Figure 5

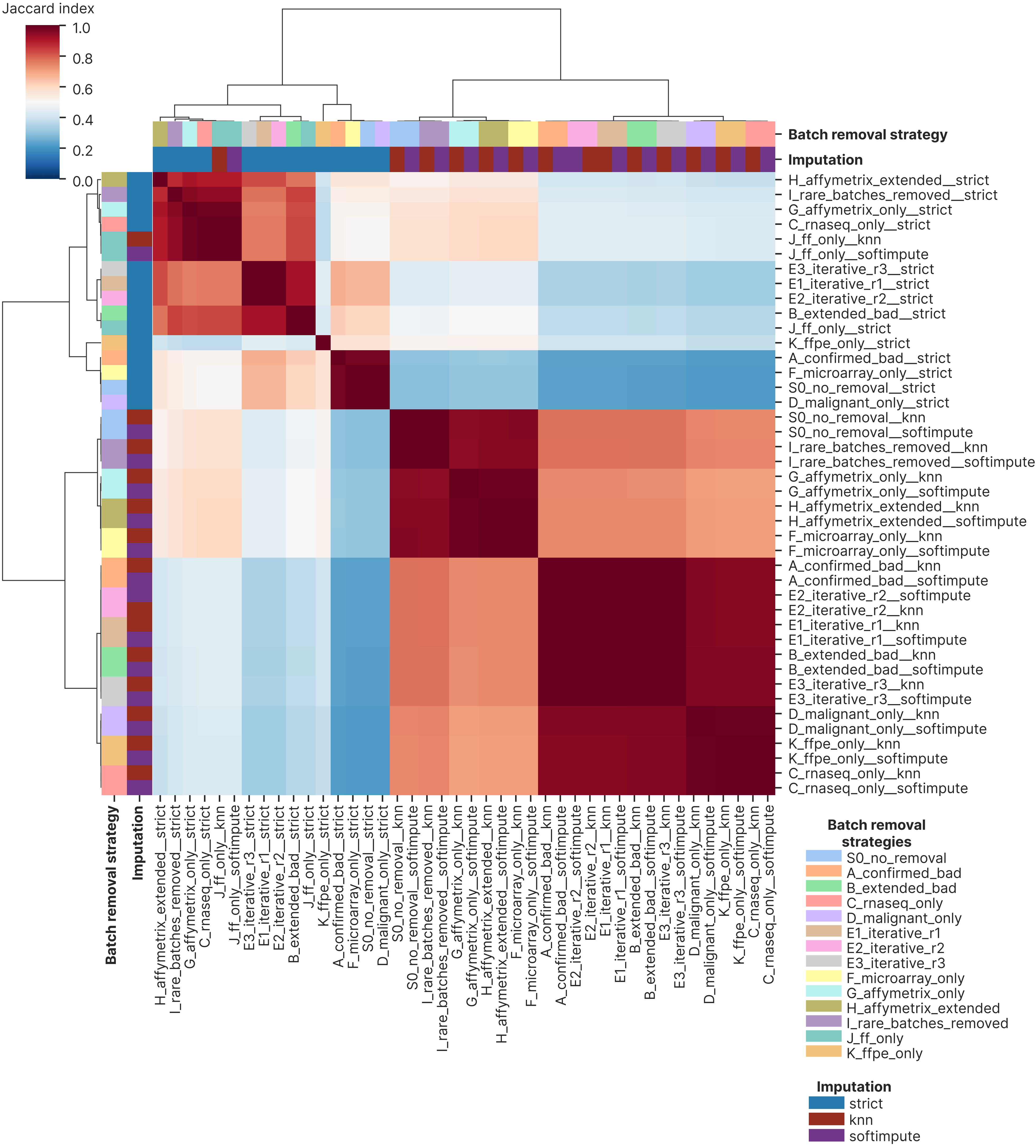

Supplementary Figure 6

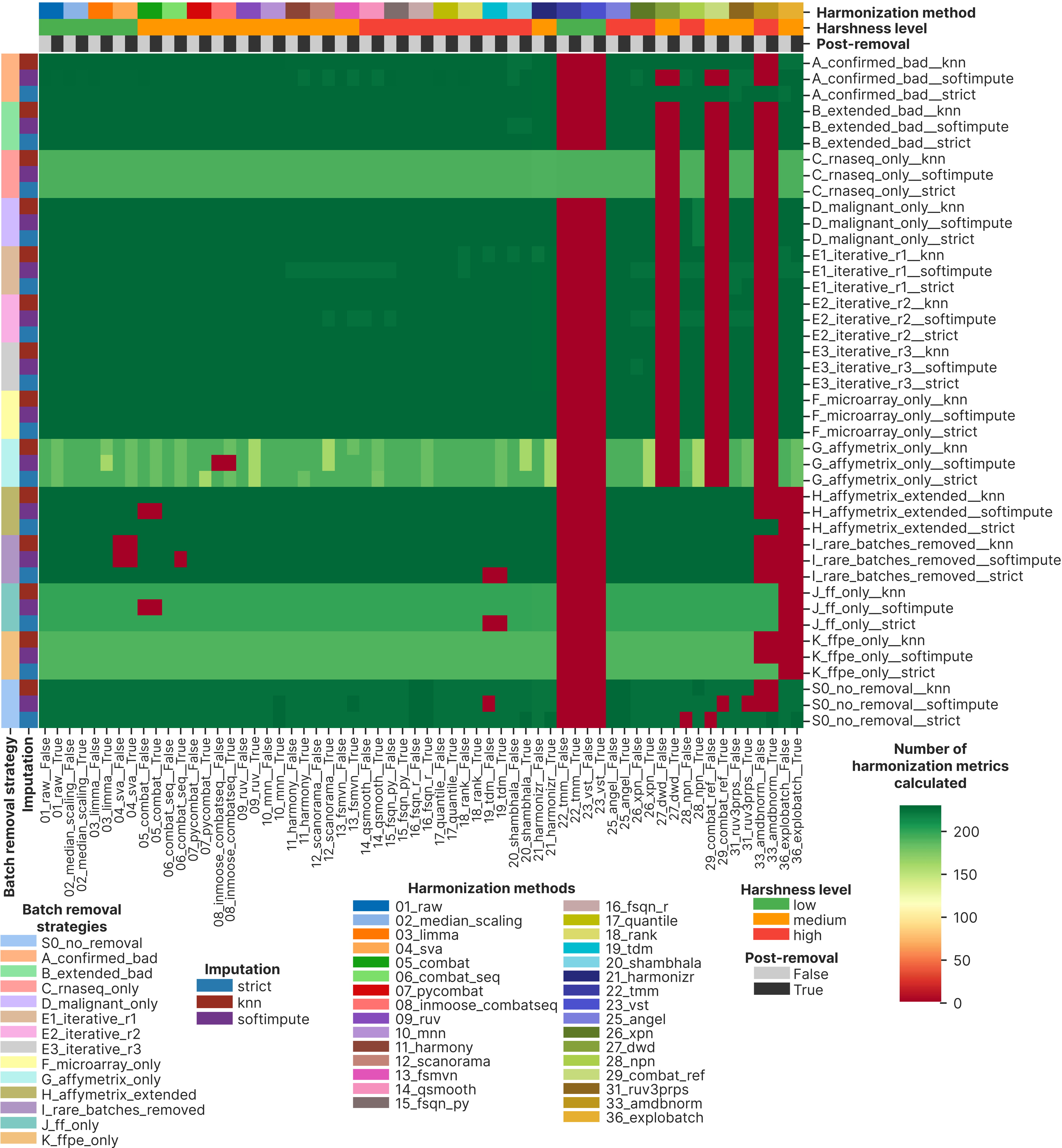

Supplementary Figure 7

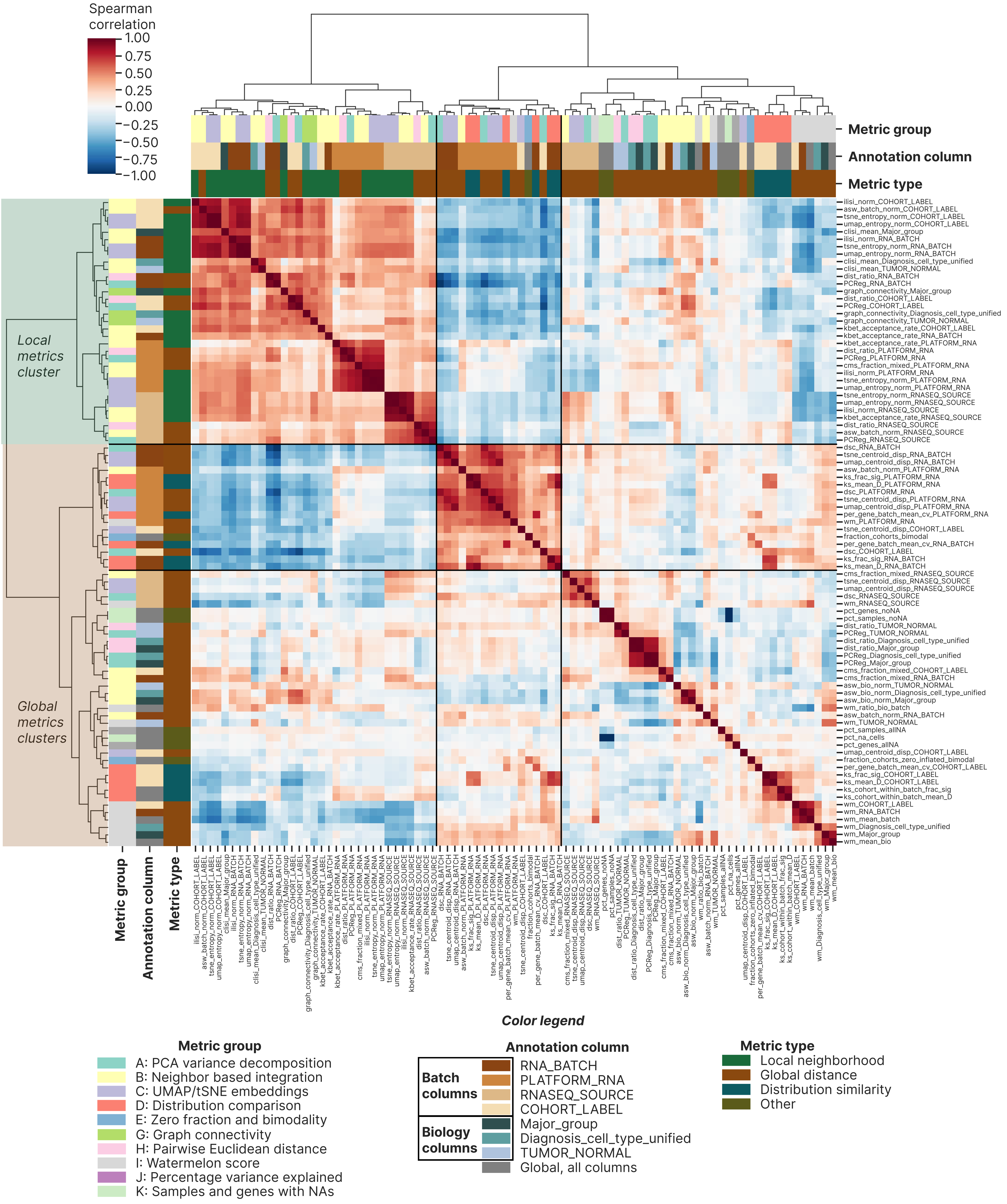

### Supplementary Figures 8-9

Supplementary Figure 8

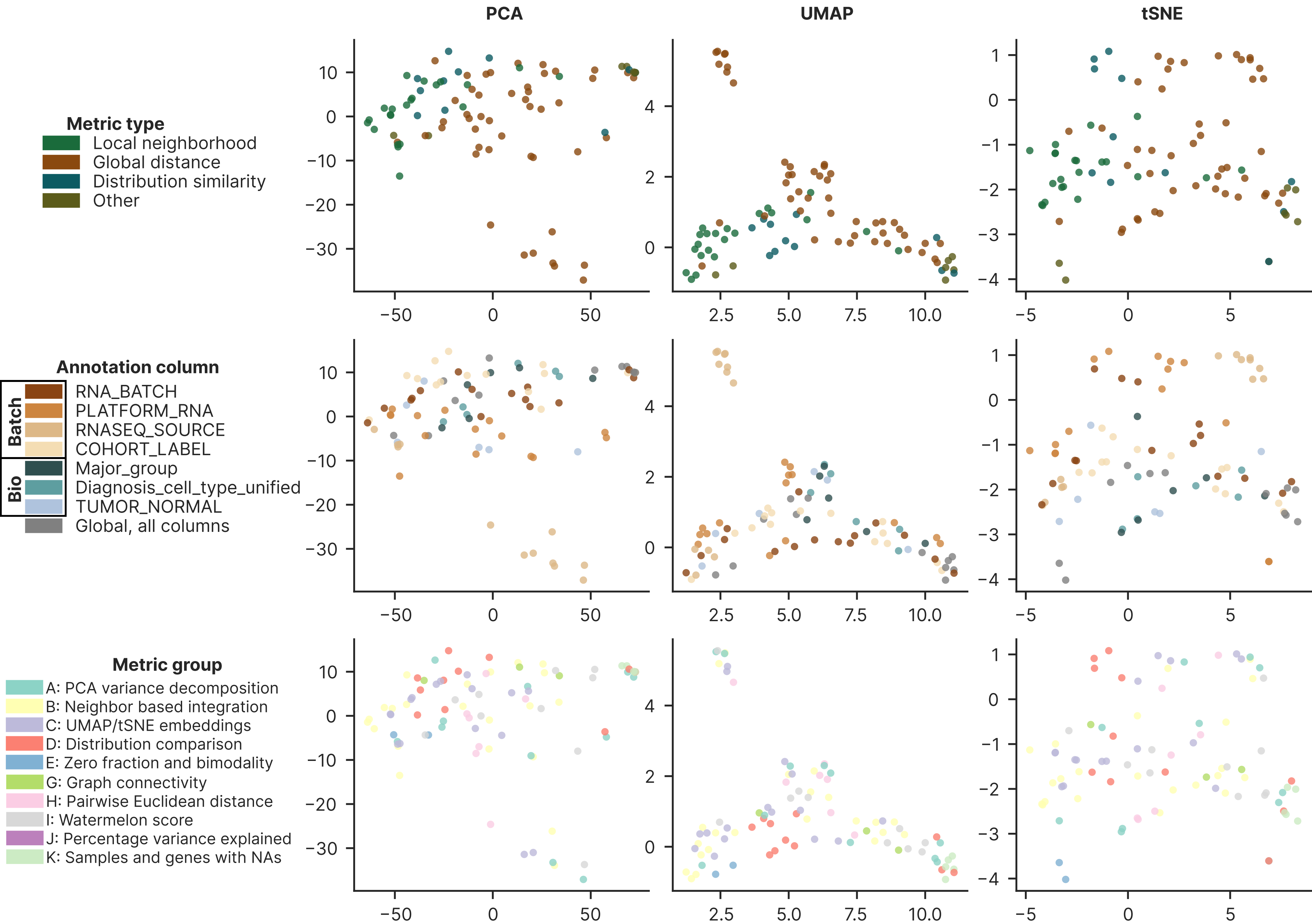

Supplementary Figure 9

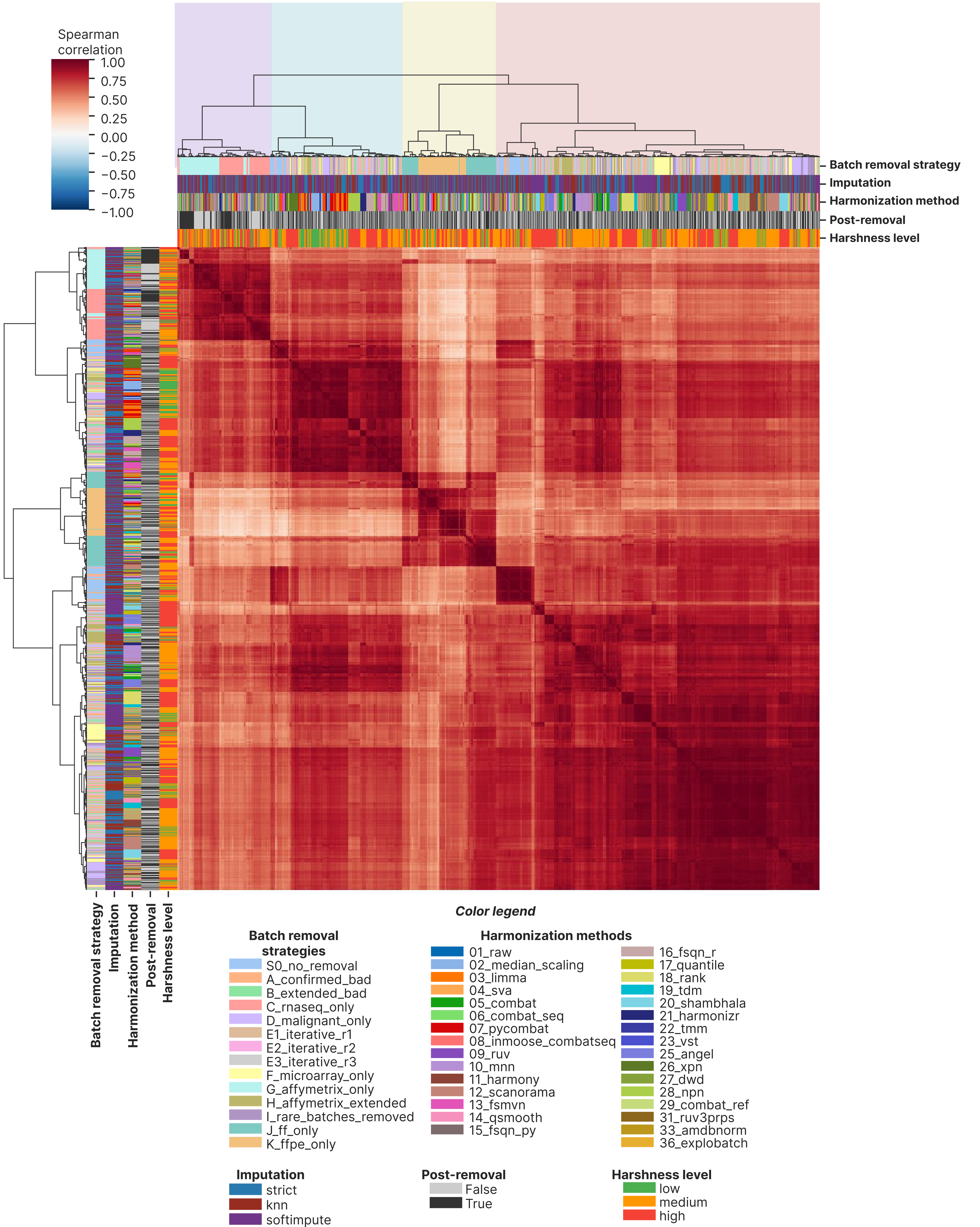

### Supplementary Figures 10-12

Supplementary Figure 10

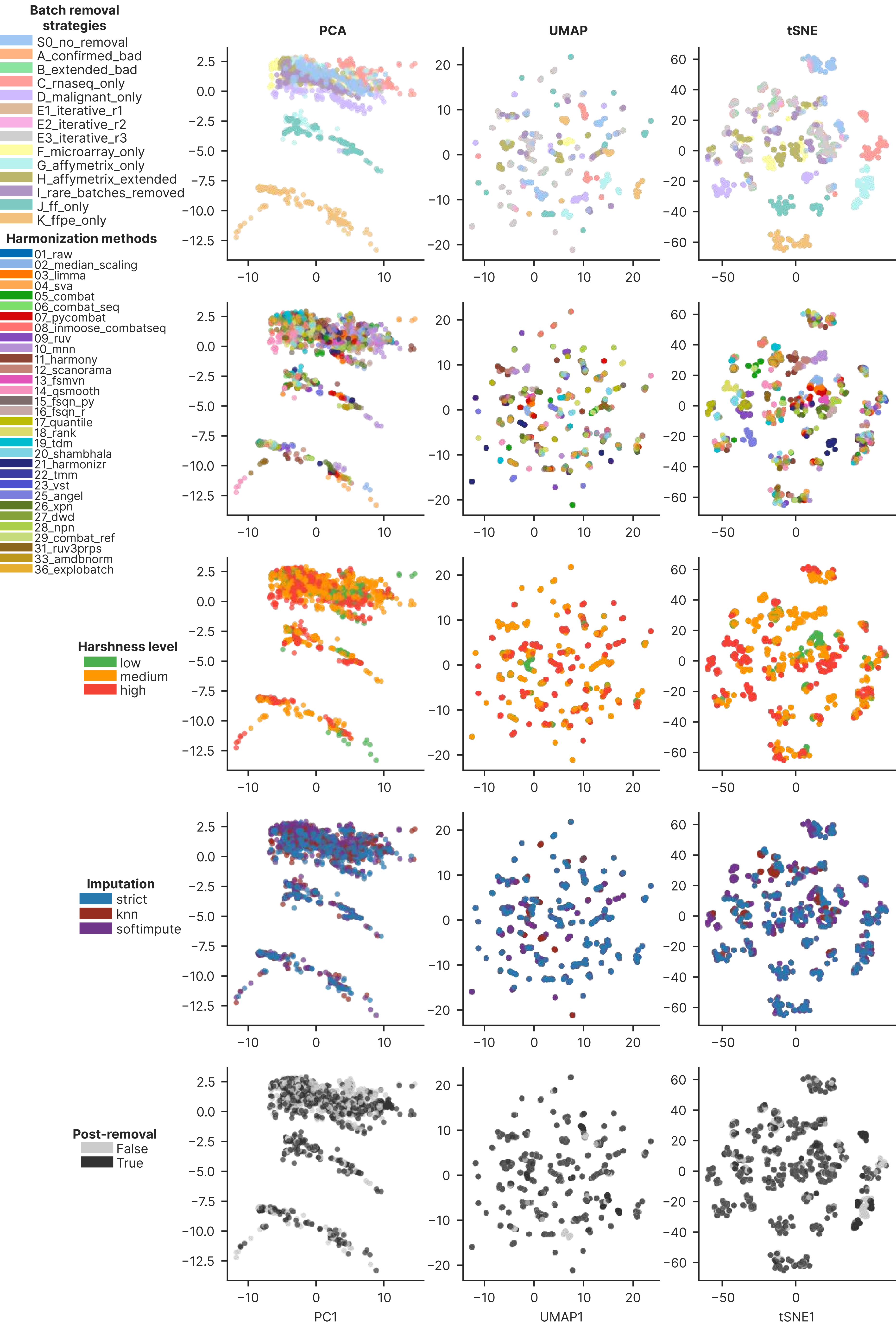

Supplementary Figure 11

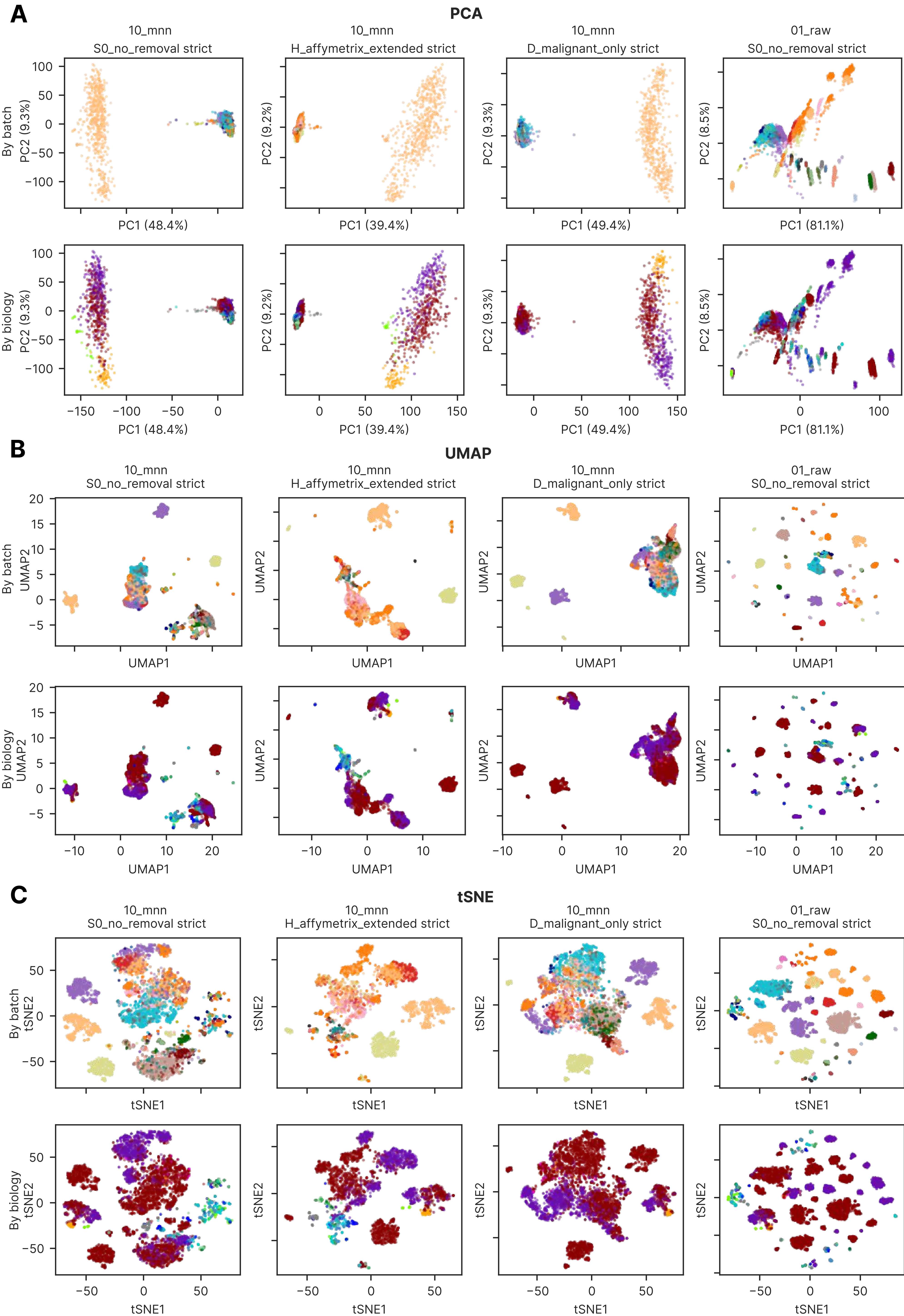

Supplementary Figure 12

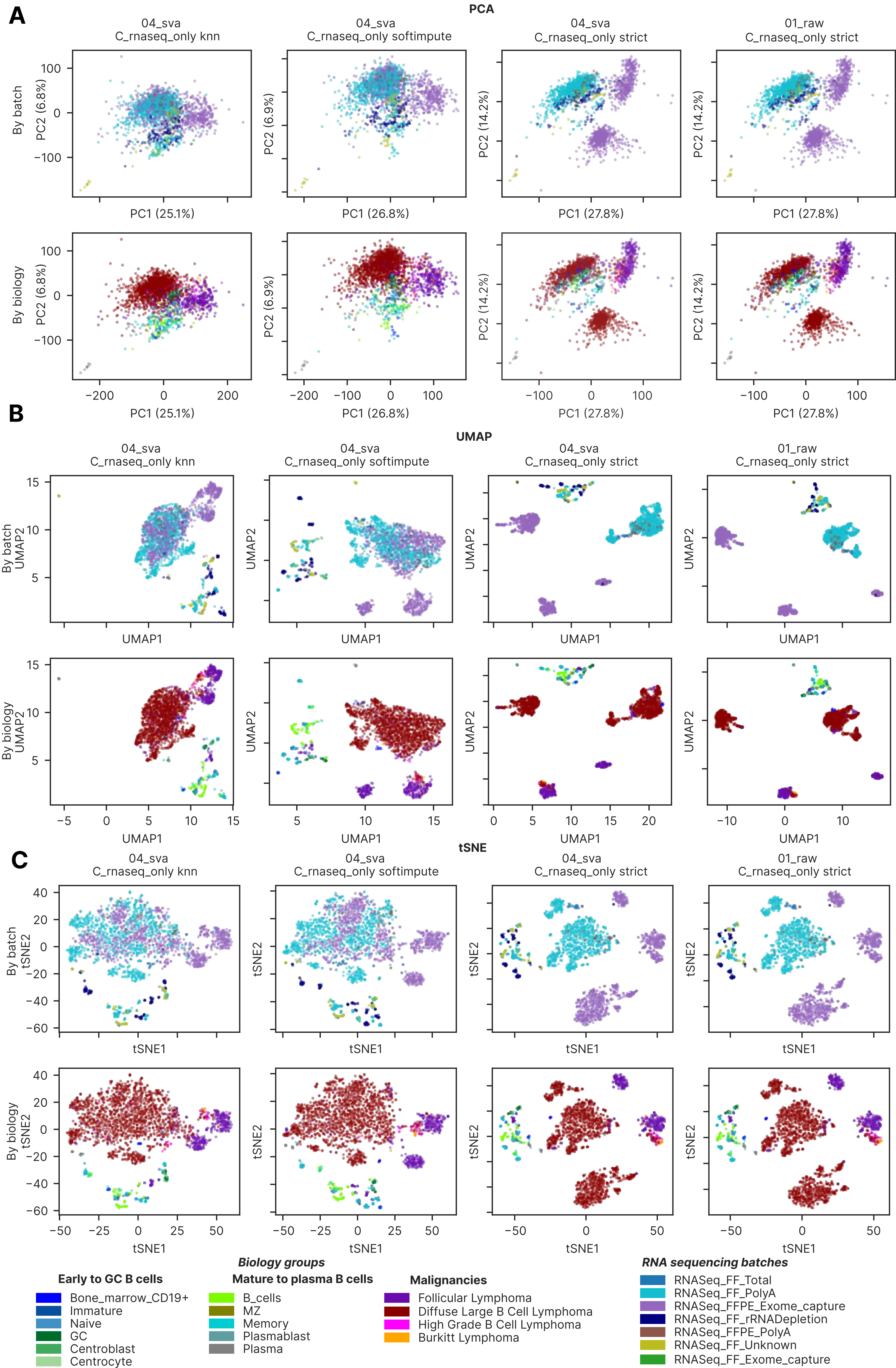

### Supplementary Figures 13-15

Supplementary Figure 13

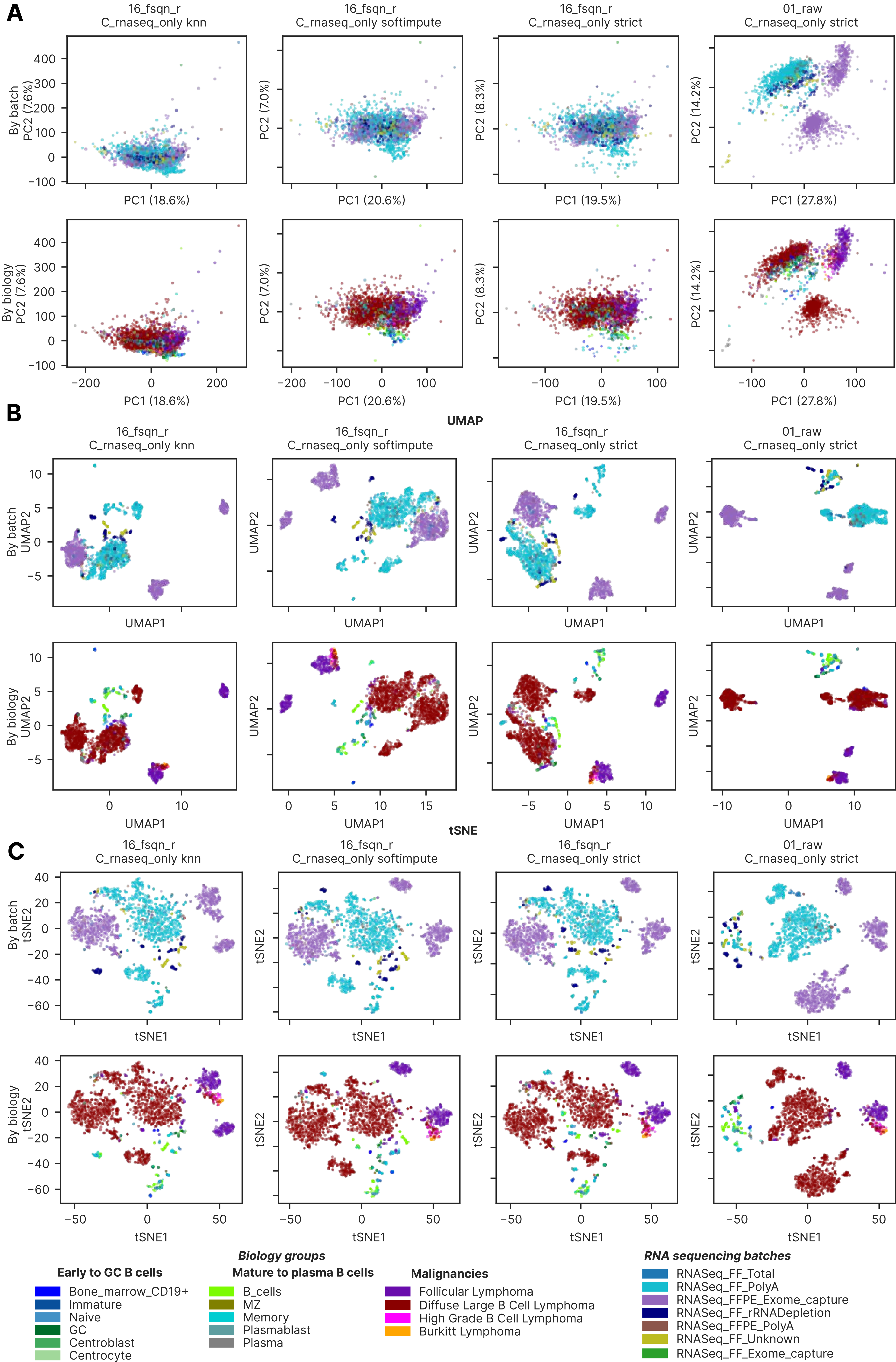

Supplementary Figure 14

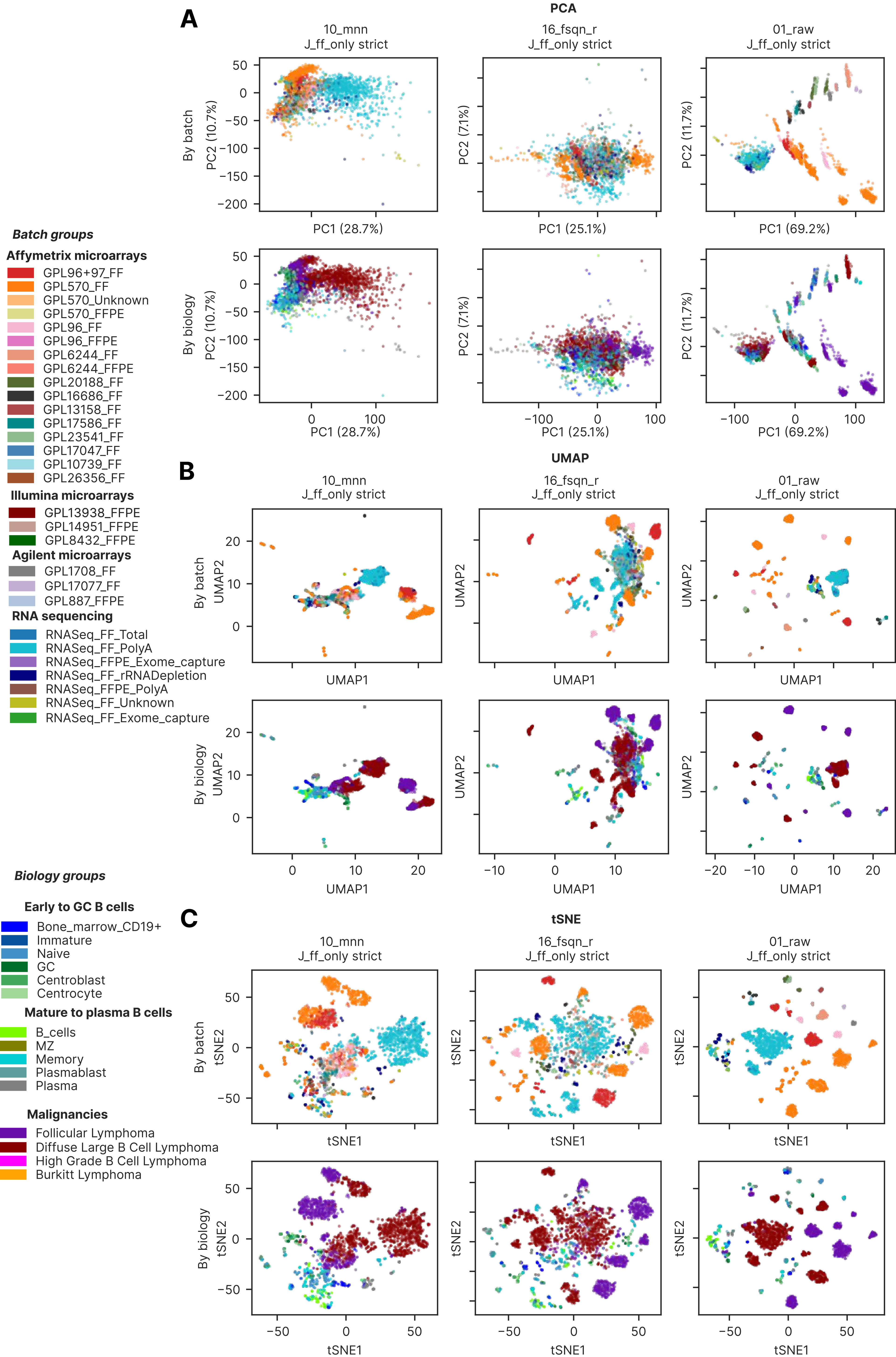

Supplementary Figure 15

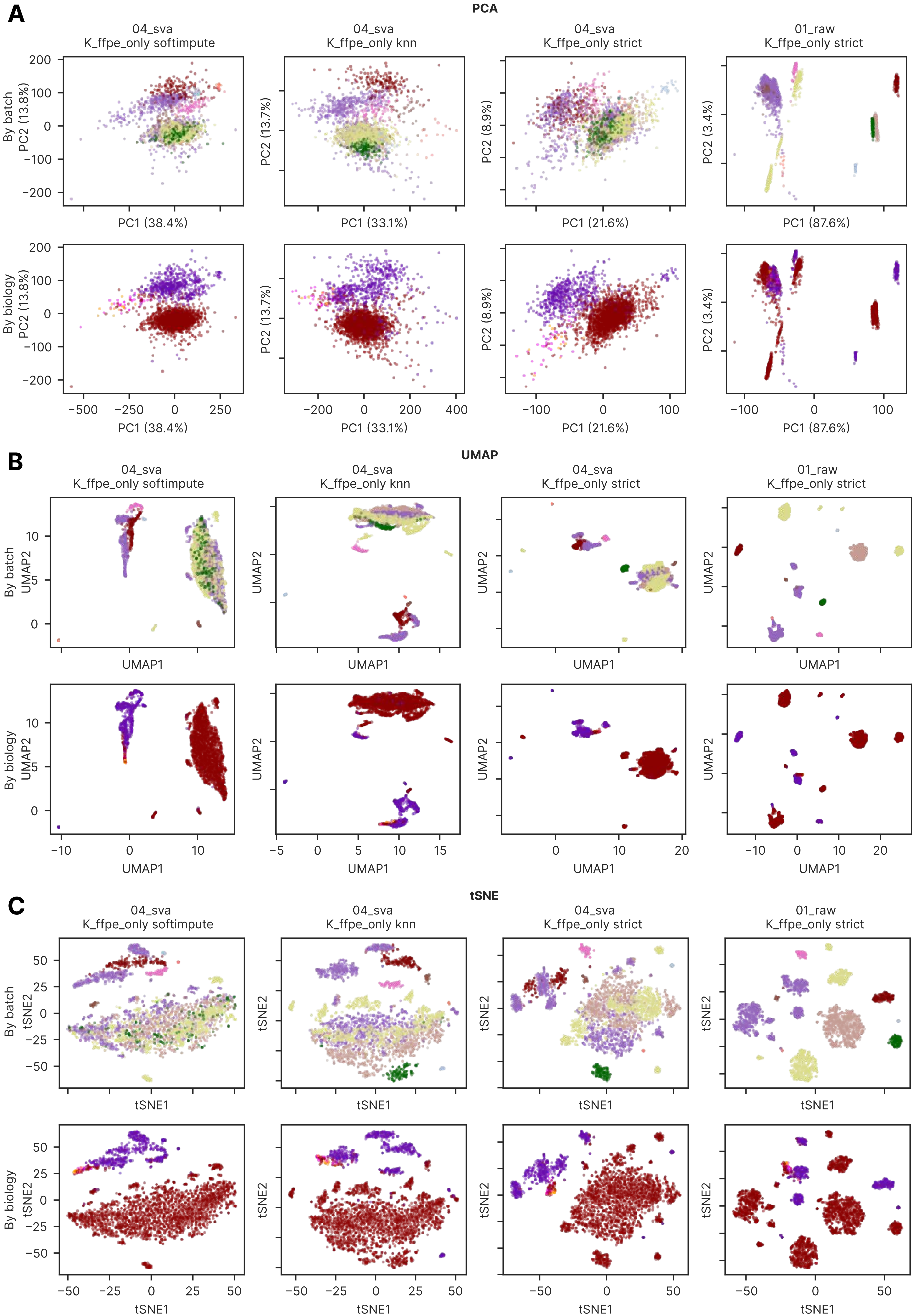

### Supplementary Figures 16-17

Supplementary Figure 16

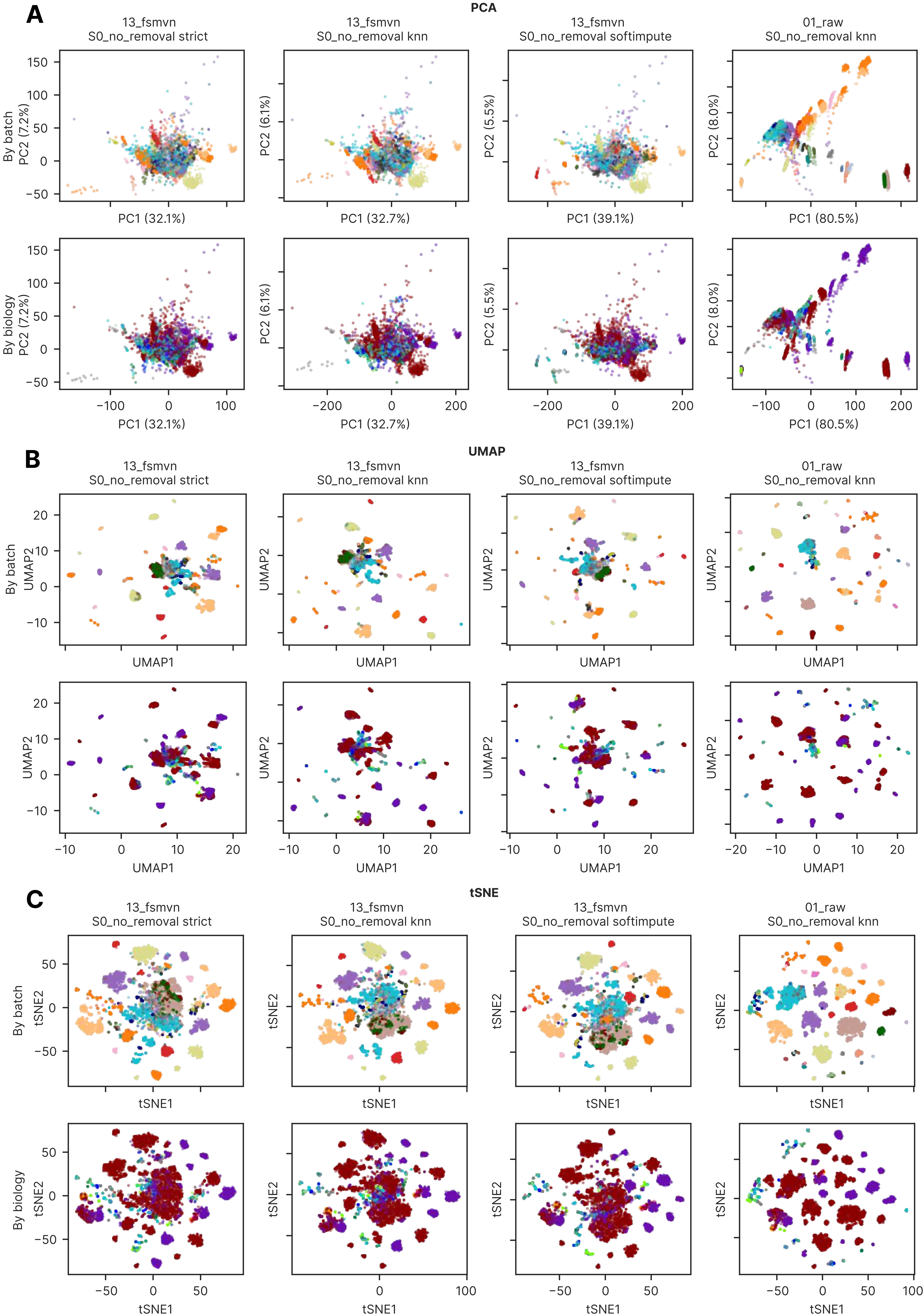

Supplementary Figure 17

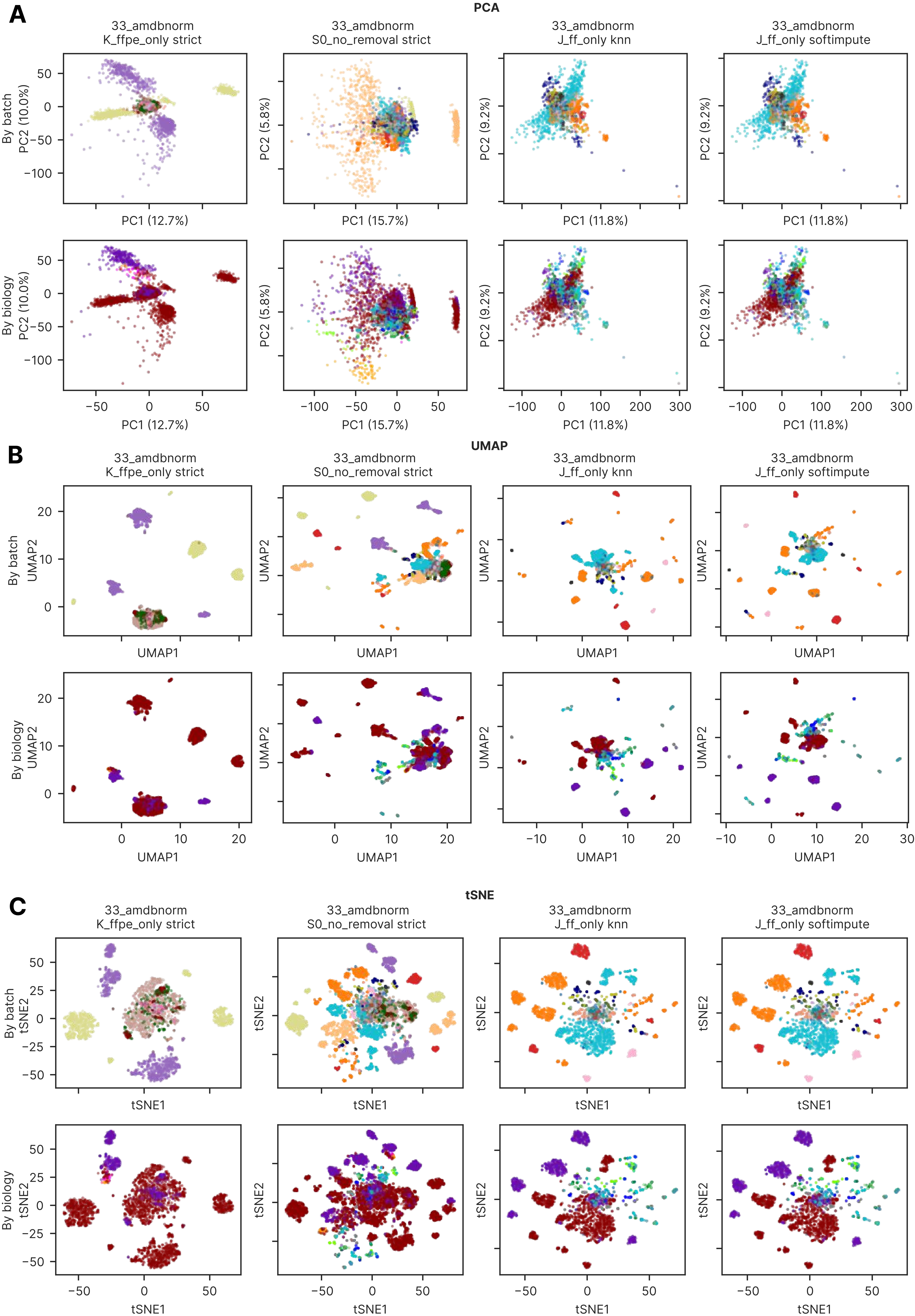
